# Discovery of a [4Fe-4S] cluster in the PRRSV Nsp1α leader protease reveals host-virus interplay in its downstream functions

**DOI:** 10.64898/2026.01.06.697956

**Authors:** Trent Quist, Anastasiya Buzuk, Henry Thanh Nguyen, Ken Takeoka, Daniel W. Bak, Eranthie Weerapana, Deborah L. Perlstein, Maria-Eirini Pandelia

## Abstract

Porcine reproductive and respiratory syndrome virus (PRRSV; *Betaarterivirus suid*) is a major global threat to swine production, yet effective antiviral therapies are lacking. The leader protease Nsp1α is essential for viral replication and innate immune suppression, and its N-terminal zinc-finger (ZF) domain is critical for function, although its molecular role remains unclear. Here, we show that the ZF domain plays only a minor role in protease activity and that Nsp1α is largely inactive following release from the polyprotein. Using Mössbauer and UV/visible spectroscopy combined with chemoproteomics, we demonstrate that the ZF site binds not only Zn but also a [4Fe-4S] cluster. Notably, the Fe-S cluster, but not Zn, allosterically modulates residual protease activity. Nsp1α directly engages the cytosolic iron-sulfur cluster assembly machinery via CIAO1 and competes with the Fe-S carrier CIAO3, establishing the [4Fe-4S] cluster as a *bona fide* cofactor. These findings redefine Nsp1α as an Fe-S-dependent viral protein and reveal new opportunities for metal-targeted antiviral strategies.

## Introduction

Porcine reproductive and respiratory syndrome virus (PRRSV; *Betaarterivirus suid*) is one of the most serious pathogens affecting swine in the United States, Europe, and China (*1, 2*). Disease control is hindered by poor vaccine efficacy and the absence of antivirals (*3–5*). PRRSV success relies heavily on its ability to suppress innate immunity early in infection, and central to this process is the nonstructural protein 1 (Nsp1), comprising two papain-like protease domains, Nsp1α and Nsp1β (*6–8*). Nsp1α is thought to be the first viral protein released from the pp1a polyprotein (*2, 9, 10*). Although its known function is to self-cleave from the polyprotein, its biological importance extends far beyond autoproteolysis: Nsp1α suppresses type I interferon responses (*7, 9, 11–13*), modulates host transcription (*12, 14*), and regulates subgenomic RNA production (*8, 15–17*), activities that are essential for viral replication, persistence, and establishing a cellular environment favorable to immune evasion. Despite this recognized multifunctionality, the molecular basis through which this small viral protease executes such diverse roles remains largely unresolved, highlighting a critical knowledge gap in our mechanistic understanding of its role in viral pathogenesis and host manipulation.

The crystal structure of Nsp1α revealed two cysteine-rich metal-binding motifs: a highly conserved N-terminal zinc finger (ZF) and a second metal-binding site located adjacent to the catalytic cysteine (*17, 18*). Both regions are essential for viral replication and immune suppression; in particular, the ZF domain governs IFN-β promoter inhibition, subgenomic RNA synthesis, and nuclear localization of Nsp1α (*11, 12, 16, 17, 19*). However, the chemical identity and mechanistic role of the bound metal ions have remained ambiguous. Although these metal sites were assigned to Zn on the basis of crystallography, the physiological relevance of Zn binding has not been defined, while Zn can inhibit thiol proteases (*20, 21*), raising questions about the true role of metal coordination in Nsp1α function.

A growing body of work challenges the assumption that cysteine-rich motifs in viral proteins invariably bind Zn and demonstrates instead that many such motifs actually bind iron-sulfur (Fe-S) clusters (*22–26*). This paradigm shift has redefined our understanding of viral metallobiology, as Fe-S cofactors have been discovered in several viral enzymes, such as the nsp12 RNA polymerase (*27*), nsp13 helicase (*28*) and the nsp14-nsp10 exoribonuclease and methyltransferase complexes from SARS-CoV-2 (*29*); the small tumor antigen of Merkel cell polyomavirus (*30*); the rotavirus nsp5 (*31*); the Hepatitis B virus X protein (HBx) (*32, 33*); and the glycine/cysteine-rich Fe-S proteins (GciS) from *Megavirinae* (*34*). While the precise functions of the Fe-S cofactors in HBx, GciS, and rotavirus nsp5 are not known, Fe-S cluster coordination enhances polymerase activity in nsp12, modulates helicase function in nsp13, enhances methyltransferase activity and regulates RNA binding in SARS-CoV-2 (*27–29*). Collectively these findings reveal a broader principle; viruses exploit the host Fe-S cluster biosynthesis machinery to acquire these metalloclusters to expand their functional repertoire, suggesting that the extent of viral Fe-S utilization has been underestimated.

Here, we identify the PRRSV Nsp1α as a *bona fide* Fe-S cluster containing protein. Using Mössbauer, UV/VIS spectroscopy, mutagenesis, and chemoproteomics, we show that Nsp1α coordinates a [4Fe-4S] cluster at its ZF site, redefining the longstanding assumption that this region functions solely as a Zn-binding domain. Although the cluster is dispensable for protease activity *in cellulo,* this activity is largely lost following cleavage of Nsp1α from the polyprotein. Fe-S coordination modulates the residual proteolytic activity in ways distinct from Zn, demonstrating that alternative metal states tune distinct biochemical functions. We further find that Nsp1α directly engages the host CIA cytosolic targeting complex (*35–37*), predominantly through CIAO1, and competes with the Fe-S carrier protein CIAO3 for access to this maturation pathway. These interactions indicate that Nsp1α relies on the host Fe-S biogenesis machinery to acquire its metallocluster and may influence Fe-S homeostasis during infection.

Although Fe-S cofactors have now been identified in several viral polymerases and helicases, no viral protease has ever been shown to coordinate or depend on an Fe-S cluster, marking Nsp1α the first of its kind. Collectively, our findings establish Nsp1α as a previously unrecognized viral Fe-S effector and expand the emerging paradigm that viruses exploit Fe-S cluster chemistry to diversify protein function. We propose that Fe-S cluster coordination in Nsp1α serves a regulatory role, to either perturb host Fe-S cluster biogenesis or modulate its immunosuppressive functions, providing a testable framework for its therapeutic targeting via selective Fe-S cluster degradation (*38*).

## Results

### The ZF motif of Nsp1α is strictly conserved across arteriviruses

To explore the evolutionary landscape of Nsp1α proteins, we generated a sequence similarity network (SSN) comprising approximately 2,200 sequences, all of which belong to the Arteriviridae family (**Fig. S1**). From this dataset, 30 species-representative sequences were selected to construct the Nsp1α phylogeny, including primate- and equine-infecting viruses as well as the distantly related Wobbly possum virus pp1a N-terminal fragment as an outgroup (*39*) (**Fig. 1A, Table S1**). Nsp1α sequences derived from simian hosts form a well-resolved group that branch separately from other mammalian lineages. The remaining sequences segregate into two major groups: an early-diverging clade containing equine arteritis virus (EAV) and a larger clade encompassing all other arteriviruses. This topology mirrors the established arterivirus phylogeny, in which Simian Hemorrhagic Fever Virus (SHFV), EAV, and Lactate Dehydrogenase Elevating Virus (LDV) precede the emergence of PRRSV genotypes 1 and 2 (*39, 40*).

**Figure 1.**
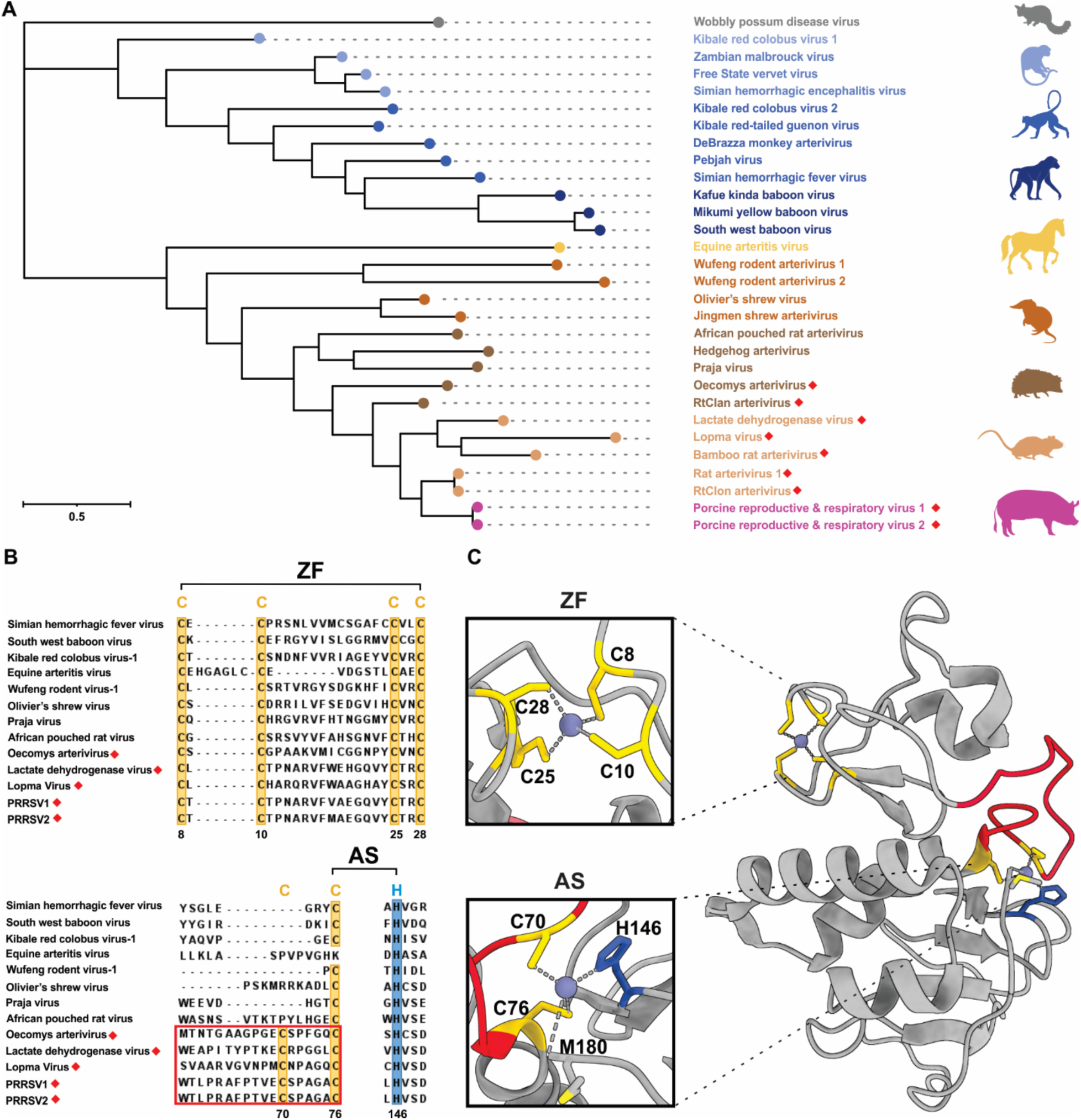
Phylogeny and multiple sequence alignment analysis of Nsp1α protein sequences from representative Arteriviruses. **A** Rooted phylogenetic tree of Nsp1α protein sequences. **B** Multiple sequence alignment of Nsp1α protein sequences from different Arterivirus species, highlighting the conserved residues involved in metal (ZF) and active site (AS) coordination. The loop insertion that contains the C70 metal binding residue is boxed in red. **C** Crystal structure of Nsp1α (PDB ID: 3IFU). Insets depict the N-terminal ZF site (top) and the proteolytic active site (AS) (bottom). The loop insertion containing C70 is colored red and the rhomb indicates the presence of C70 in the arteriviral sequence.

To investigate any functional diversification across Nsp1α orthologs, we next generated a multiple sequence alignment of representative Arterivirus proteins (**Fig. 1B, Fig. S2**). The N-terminal zinc finger (ZF) motif, defined by four invariant cysteine residues, is strictly conserved across all sequences, underscoring its fundamental importance as a shared structural and functional element of Nsp1α. In contrast, the catalytic dyad (C76 and H146, PRRSV numbering) is conserved in all arteriviruses except EAV, in which Nsp1 remains uncleaved due to the absence of the catalytic cysteine (**Fig. 1B**) (*11, 17*). Although proteolytic processing of Nsp1α in SHFV has previously been debated, the preservation of the catalytic dyad in our alignment supports more recent evidence that SHFV Nsp1α undergoes cleavage (*11, 41*). Thus, while protease activity is broadly retained across the family, it is not universally required. By contrast, the absolute conservation of the ZF motif across all arteriviruses points to a non-redundant and central role in Nsp1α-mediated host-virus interactions.

We further examined the conservation of a putative second Zn-binding site by analyzing the presence of cysteine C70, which together with the carboxy-terminal of M180 has been proposed to coordinate a catalytic Zn ion in the available crystal structure (**Fig. 1C)** (*18, 42, 43*). C70 is absent from early diverging Nsp1α proteins in primate-, shrew-, and equine-infecting viruses and is exclusively found in rodent-porcine lineages, indicating that this feature arose later in evolution (**Fig. 1A, Fig. S2**). Its localization within a lineage-specific loop insertion further suggests that this auxiliary metal-binding site represents a specialized adaptation rather than a core functional element. Collectively, these analyses establish the N-terminal ZF motif as the only universally conserved structural feature of Nsp1α, implicating it as a primary determinant of Nsp1α function across the Arteriviridae family.

### Nsp1α binds an Fe-S cluster in its N-terminal ZF domain

Heterologous expression of wild type Nsp1α resulted in its accumulation in inclusion bodies that when solubilized in 8 M urea produced a reddish-brown solution (**Fig. S3**). The color persisted under O_2_-free conditions following affinity purification under denaturing conditions. The 80 K Mössbauer spectrum of this sample is characterized by a quadrupole doublet with δ = 0.26 mm/s and ΔE_Q_ = 0.49 mm/s, parameters characteristic of high-spin Fe(III) ions in [2Fe-2S]^2+^ clusters (**Fig. 2A**). This assignment was further supported by the optical spectrum of this sample, which exhibited bands at 325 nm, 419 nm, and 460 nm, characteristic of tetracysteine-ligated [2Fe-2S] clusters (**Fig. 2B**) (*33, 44*).

**Figure 2.**
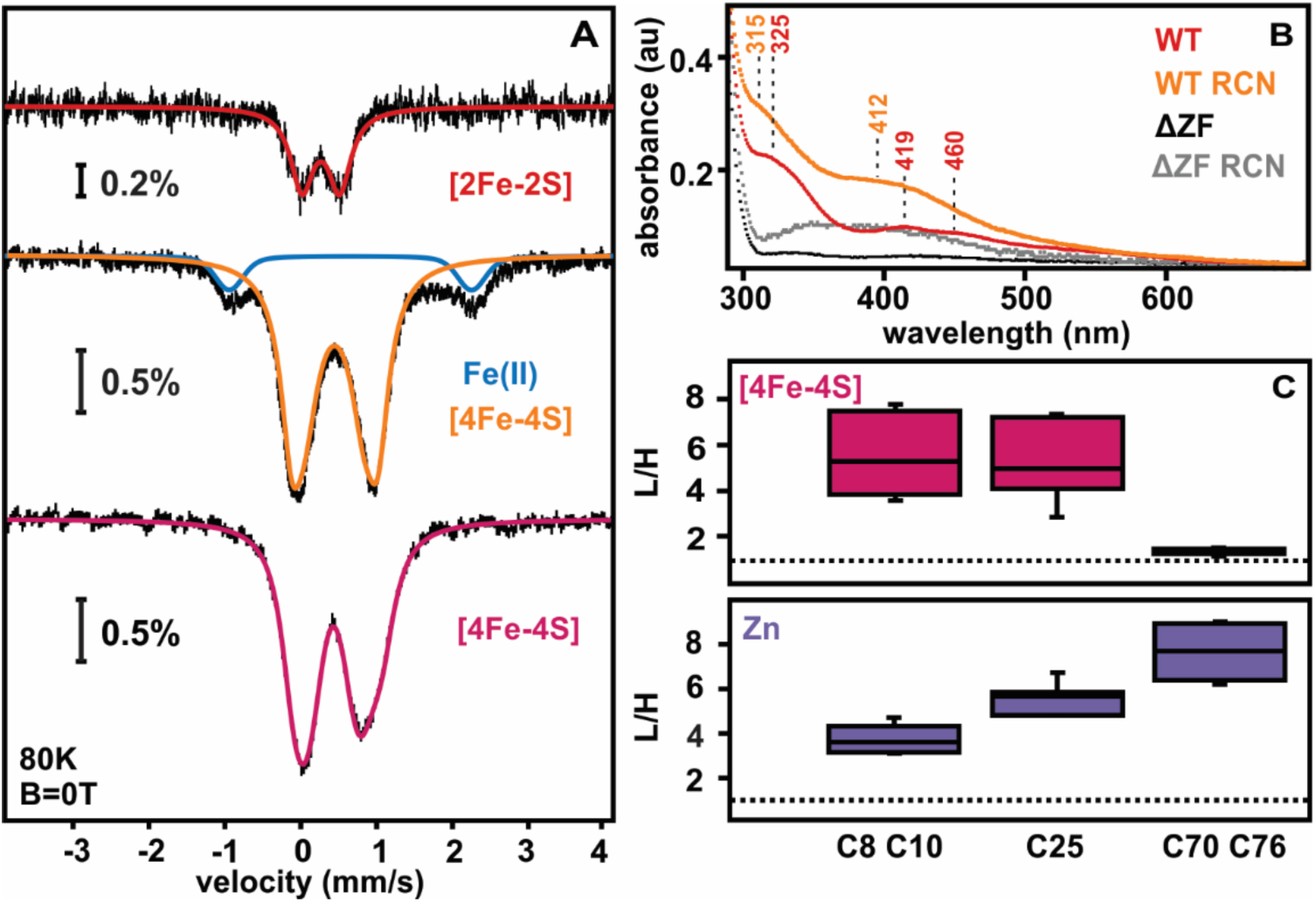
Nsp1α binds an Fe-S cluster at the N-terminal ZF site. **A** Mössbauer spectra of the Nsp1α^WT^ protein purified in urea from inclusion bodies as isolated (top) and after chemical reconstitution with Fe(II) and sulfide under O_2_-free conditions (middle). Mössbauer spectra of natively purified, semi-enzymatically reconstituted Nsp1α (bottom). **B** Optical spectra of wild type Nsp1α and Nsp1α^ιΔZF^ purified from inclusion bodies as isolated (red and black traces, respectively) and after chemical reconstitution (orange and gray traces, respectively). **C** Chemoproteomic cysteine reactivity analysis of natively purified Nsp1α reconstituted with either Fe(II) and sulfide (magenta) or Zn (purple). Quantification of cysteine reactivty was achieved through isotopic NEM-labeling (light -Nsp1α^apo^; heavy - Nsp1α^Fe-S^ or Nsp1α^Zn^) and subsequent LC-MS/MS analysis. Light:heavy (L/H) ratios are displayed as box plots and the median L/H ratio value across all cysteine is indicated by a dashed line.

To address whether the [2Fe-2S] cluster is an oxidative byproduct or a true cofactor, we performed an additional chemical reconstitution, similarly followed by affinity purification under denaturing conditions. The 80 K Mössbauer spectrum of the chemically reconstituted Nsp1α is now dominated by a quadrupole doublet with δ = 0.45 mm/s and ΔE_Q_ = 0.95 mm/s, parameters characteristic of [4Fe-4S]^2+^ clusters (**Fig. 2A**) (*33, 45*). This assignment is further corroborated by the different optical spectra that are now dominated by a broad absorption band at 412 nm. Identical experiments were performed with the Nsp1α^ΔZF^ variant, in which all 4 cysteines are replaced by alanines. Nsp1α^ΔZF^ showed no optical features in the region associated with Fe-S clusters. Chemical reconstitution failed to incorporate an Fe-S cluster as indicated both by the absence of color in the elution fractions and lack of any Fe-S cluster optical features, confirming that cluster binding depends on the four N-terminal cysteines (**Fig. 2B**).

We also obtained Nsp1α from expression of the self-cleaving Nsp1αβ (Nsp1^WT^) polyprotein that releases soluble Nsp1α (under native conditions), which, however, does not purify with any metals when grown in minimal media as determined by ICP-AES (**Table S2**). We employed the cysteine desulfurase IscS to perform a semi-enzymatic Fe-S cluster reconstitution to generate the soluble cofactor-enriched Nsp1α (*46*). The Mössbauer spectrum of the semi-enzymatically reconstituted Nsp1α confirms incorporation of a [4Fe-4S]^2+^ cluster, showing a broader quadrupole doublet best fit by two overlapping doublets with average parameters δ = 0.43 mm/s and ΔE_Q_ = 0.86 mm/s (**Fig. 2A**). These values, despite slight differences owing to sample heterogeneity between denaturing and native conditions, consistently support the assignment of the Nsp1α cofactor as a [4Fe-4S] cluster, with the [2Fe-2S] form likely generated as an artifact of misassembly or degradation.

EPR spectroscopy showed that the [4Fe-4S]^2+^ cluster of Nsp1α could not be reduced to the 1+ state with sodium dithionite nor oxidized to the 3+ state with potassium ferricyanide, aside from oxidative degradation to a [3Fe-4S]^1+^ cluster (*33, 45*)(**Fig. S4**). These results demonstrate that the Fe-S cluster does not have an active redox role and is prone to oxidative degradation. We examined the oxygen sensitivity of the [4Fe-4S]^2+^ cluster in Nsp1α by stopped-flow absorption experiments and confirmed that the cluster rapidly decomposes with an average rate of 0.042 min^-^_1_, as demonstrated by the progressive loss of the band at 412 nm (**Fig. S5**).

To complement our spectroscopic analyses, we next probed the metal-binding specificity of Nsp1α using a quantitative chemoproteomics strategy (*47*). This approach exploits differential thiol reactivity to identify cysteine residues protected by metal binding. Apo (cofactor devoid) and holo (cofactor enriched) forms of Nsp1α were selectively alkylated with isotopically distinct variants of N-ethylmaleimide (NEM-d^0^, light, and NEM-d^5^, heavy, respectively), followed by proteolytic digestion and quantitative LC–MS/MS analysis. Because metal coordination shields cysteines from alkylation, cysteines involved in metal binding exhibit elevated light-to-heavy (L/H) peptide ratios relative to non-coordinating residues. Reduced cysteine reactivity was observed at both the N-terminal ZF cysteines (C8, C10, and C25) and the active site cysteines (C70 and C76), indicating that Zn^2+^ can associate with both regions of the protein (**Fig. 2C**). In contrast, reconstitution with a [4Fe-4S]^2+^ cluster resulted in selective protection of the ZF cysteines (C8, C10, and C25), while the active-site cysteines remained fully accessible to NEM labeling (**Fig. 2C**). This pattern indicates preferential and exclusive coordination of the Fe-S cluster at the N-terminal ZF motif.

To further quantify the metal-binding preferences of Nsp1α, we measured Zn^2+^ binding by isothermal titration calorimetry. These experiments revealed that the ZF motif binds Zn^2+^ with moderate affinity (Kd ∼ 1 μM), whereas the active site displays negligible Zn^2+^ binding under identical conditions and in the absence of the ZF site (**Fig. S6**). Together, these data demonstrate that although Zn^2+^ can associate with multiple cysteine-rich regions of Nsp1α, the [4Fe-4S] cluster is selectively ligated at the N-terminal ZF, establishing this motif as the exclusive physiological site for Fe-S cluster coordination.

### Nsp1α is largely inactive post-cleavage from Nsp1

To determine whether the metallocofactor identity modulates Nsp1α protease function, we first examined Nsp1 *in cis* autoproteolysis in a cellular context. The full-length Nsp1^WT^ was expressed under conditions that selectively favored incorporation of Fe, Zn, both metals, or neither, and proteolytic processing was assessed by SDS-PAGE and Western Blotting. In all conditions tested, Nsp1^WT^ underwent efficient cleavage into Nsp1α and Nsp1β (**Fig. 3A**), indicating that self-processing is robust and insensitive to the identity or availability of metal cofactors. Consistent with this observation, Nsp1 variants lacking the N-terminal ZF site (ΔZF) or the auxiliary cysteine C70 also retained normal *cis*-autoproteolytic activity. In contrast, mutation of the catalytic nucleophile (C76A) or the general base (H146A) abolished cleavage (**Fig. 3B**), confirming that processing arises from intrinsic Nsp1α protease activity rather than host proteases. Together, these results establish that metallocofactor binding is not required for Nsp1 *cis*-autoproteolysis.

**Figure 3.**
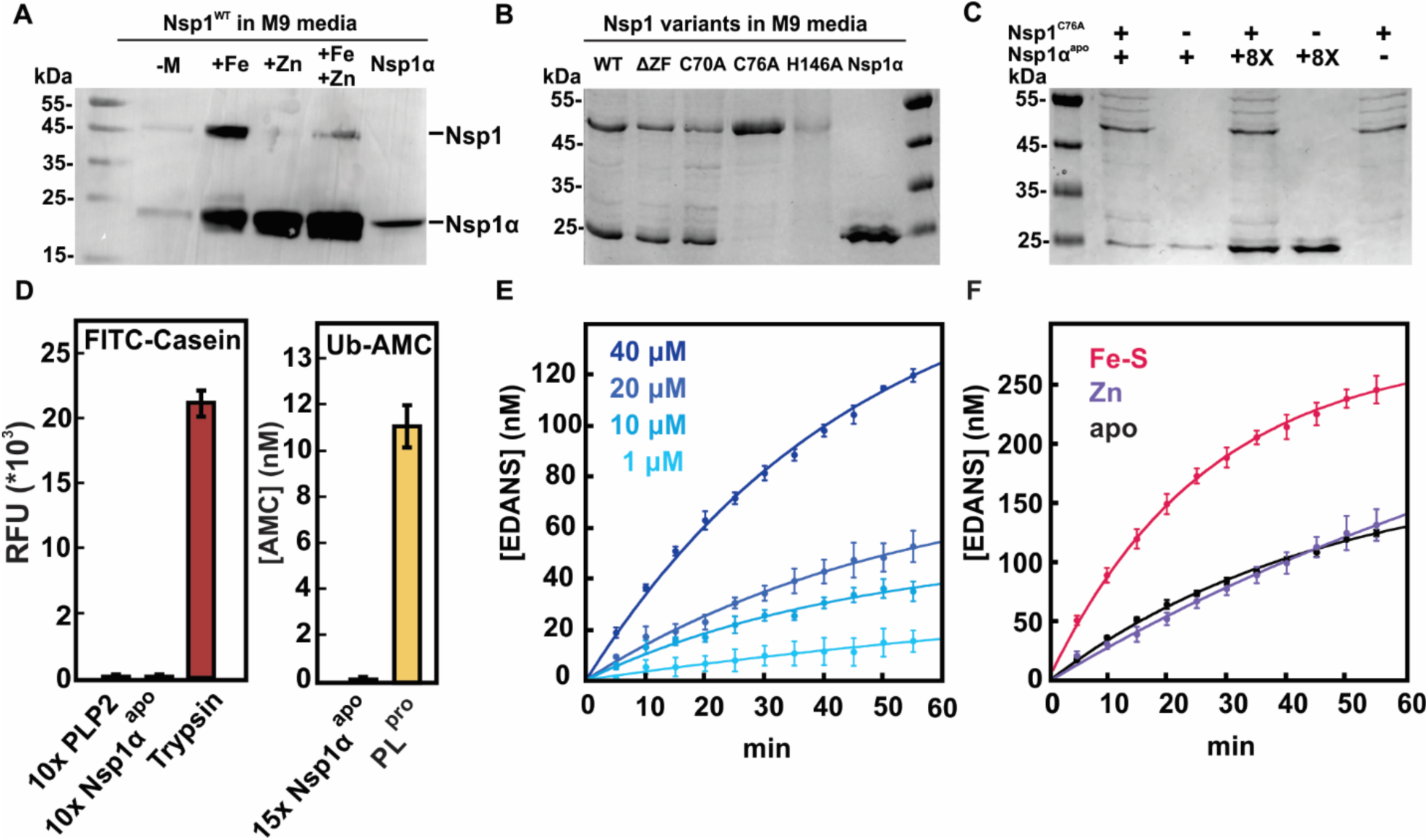
Nsp1α protease activity against a variety of substrates. **A** Western blot of *E. coli* lysates expressing Nsp1^WT^ in minimal media excluding metals (-M) or supplemented with Fe and or Zn. **B** SDS-PAGE of *E. coli* lysates grown in minimal media supplemented with Fe expressing either Nsp1^WT^, Nsp1^ΔZF^, Nsp1^C70A^, Nsp1^C76A^ or Nsp1^H146A^. **C** SDS-PAGE of Nsp1α^apo^ *in vitro* protease assay against the full-length Nsp1 polyprotein substrate. **D** Nsp1α^apo^, and PLP2 protease activity against fluorescent substrate FITC-casein compared with that of trypsin (left). Deubiquitinase activity of Nsp1α againats Ub-AMC compared to that of PL^pro^ (right). **E** Nsp1α^apo^ dose-dependent *in vitro* protease activity against the FECAMATVYD peptide substrate. **F** Nsp1α^Fe-S^ vs Nsp1α^apo^ vs Nsp1α^Zn^ protease activity against the FECAMATVYD peptide substrate.

We next asked whether Nsp1α retains proteolytic activity once liberated from the polyprotein and whether this activity could be modulated by metal binding. To address this, we reconstituted protease activity using purified components *in vitro*, initially focusing on metal-free Nsp1α (Nsp1α^apo^) to exclude contributions from co-purifying metals. A catalytically inactive Nsp1^C76A^ variant was also generated and validated for use as a substrate (**Fig. S7**). Because full-length Nsp1 remains poorly soluble in isolation, we tested several constructs designed to stabilize the substrate, including fusion to a region of the adjacent Nsp2 sequence and MBP tagging (*48*). While these approaches yielded soluble protein, Nsp1α^apo^ did not cleave Nsp1^C76A^ *in trans* under any condition tested, even at elevated enzyme concentrations or varying pH (**Fig. 3C, Fig. S8**). In addition, in the Nsp1^C76A^-Nsp2 (aa:1-501) construct, Nsp1β remained proteolytically active in the lysates (**Fig. S7**) regardless of the presence of the N-terminal inactive Nsp1α ^C76A^ fragment. This result indicates that downstream proteolytic processing does not require prior release of Nsp1α, in agreement with previous reports (*7, 8*).

To assess whether Nsp1α possesses broader proteolytic activity that might be masked by substrate architecture, we next evaluated its activity against generic and tailored fluorogenic substrates. Nsp1α exhibited minimal activity toward FITC-labeled casein, comparable to that observed for the paralog PRRSV PLP2 protease under identical conditions and orders of magnitude lower than trypsin (**Fig. 3D**) (*49*). Given that the PLP2 protease also functions as a deubiquitinase, we further tested Nsp1α against Ub-AMC and the canonical deubiquitination substrate Z-RLRGG-AMC (*50*). In both cases, Nsp1α showed no detectable activity, whereas PRRSV PLP2 as well as the SARS-CoV PL^pro^ efficiently cleaved these substrates (**Fig. 3D, Fig. S9**) (*49, 51*).

Finally, to probe sequence-specific proteolysis, we designed a fluorogenic peptide encompassing the native Nsp1 cleavage motif (FECAMATVYD) flanked by a DABCYL-EDANS FRET pair (*18, 52*). Proteolytic cleavage of this peptide will separate fluorophore and quencher, resulting in increased fluorescence. Although Nsp1α^apo^ catalyzed cleavage of this substrate at a low but reproducible rate (∼10^-3^ min^-1^) (**Fig. 3E**), the observed activity was dose-dependent and abolished by the C76A mutation, further demonstrating specificity. Nevertheless, the catalytic efficiency was markedly lower than that of trypsin, even at substantially higher Nsp1α^apo^ concentrations (**Fig. S10**), indicating that Nsp1α is either intrinsically a slow protease or becomes autoinhibited following release from the polyprotein. Together with the minimal activity of both PRRSV Nsp1α and PLP2 toward a generic protease substrate (FITC-casein), these findings support a model in which Nsp1α (and PLP2) function predominantly *in cis* during polyprotein processing.

Although the activity against the peptide substrate is low, we sought to utilize this assay to assess the impact of metal ions on Nsp1α, for which proteolytic activity remains the only established enzymatic function. Considering that Nsp1α contains two Zn ions in its crystal structure (*18*), we first explored whether Zn acts as a positive or negative effector of protease activity. Zn had no measurable effect on Nsp1α activity at stoichiometric concentrations; however, at excess concentrations, it inhibited activity (**Fig. S10**). In contrast, reconstitution of Nsp1α with a [4Fe-4S]^2+^ cluster resulted in an approximately fourfold increase in peptide cleavage relative to the apo protein (**Fig. 3F**). Although modest in magnitude, this enhancement was reproducible and specific, indicating that Fe-S cluster binding exerts a positive allosteric effect on the protease active site despite being spatially separated from the N-terminal ZF site. Thus, Fe-S cluster binding modestly enhances proteolytic activity, suggesting a potential regulatory link between metal incorporation and enzymatic function, which was not discernible in our *in cellulo* experiments. Together, these findings establish that while Nsp1α proteolysis is not strictly metal-dependent, Fe-S cluster incorporation can tune enzymatic activity through allosteric coupling, consistent with a model in which metal binding serves a modulatory role.

### Nsp1α recruits the cytosolic Fe-S assembly targeting complex

Viruses are not known to encode the machineries required for maturation of Fe–S proteins, making the presence of a [4Fe-4S] cluster in the arteriviral Nsp1α especially intriguing. Because Fe-S clusters cannot self-assemble *in vivo*, any viral Fe-S protein must access the host’s cluster biogenesis pathway, raising the fundamental question of how Nsp1α acquires its metallocluster during infection. If the cluster in Nsp1α is biologically relevant, the protein should be able to engage one or more of the cytosolic iron-sulfur cluster assembly (CIA) factors as this system is responsible for installing Fe-S cofactors into eukaryotic cytosolic and nuclear proteins (*27–29, 37*).

To test this prediction and assess the biological relevance of the [4Fe-4S] cluster in Nsp1α, we investigated whether the protein engages with components of the CIA machinery responsible for targeting and delivering clusters to recipient proteins. Nsp1α reproducibly co-eluted with the complete CIA targeting complex (CTC; Cia1-Cia2-Met18) (*37, 53*), which identifies apo-proteins and inserts their metallocluster (**Fig. 4A**). Nsp1α co-eluted with Cia1 and Cia2 irrespective of the presence of Met18, demonstrating that interaction with the *Sc*CTC is primarily mediated through Cia1-Cia2 and does not require Met18. In contrast, a weaker recovery of Nsp1α in the eluate was observed with the early-acting CIA scaffold complex (Nbp35-Cfd1, **Fig. S4B, Fig. S11**) (*37*), and minimal binding was observed with an unrelated control protein, PPAR-γ (**Fig. S12**) (*54*). These results indicate that Nsp1α is recognized by the CTC, supporting the hypothesis that it receives its Fe-S cofactor from the host biogenesis machinery.

**Figure 4.**
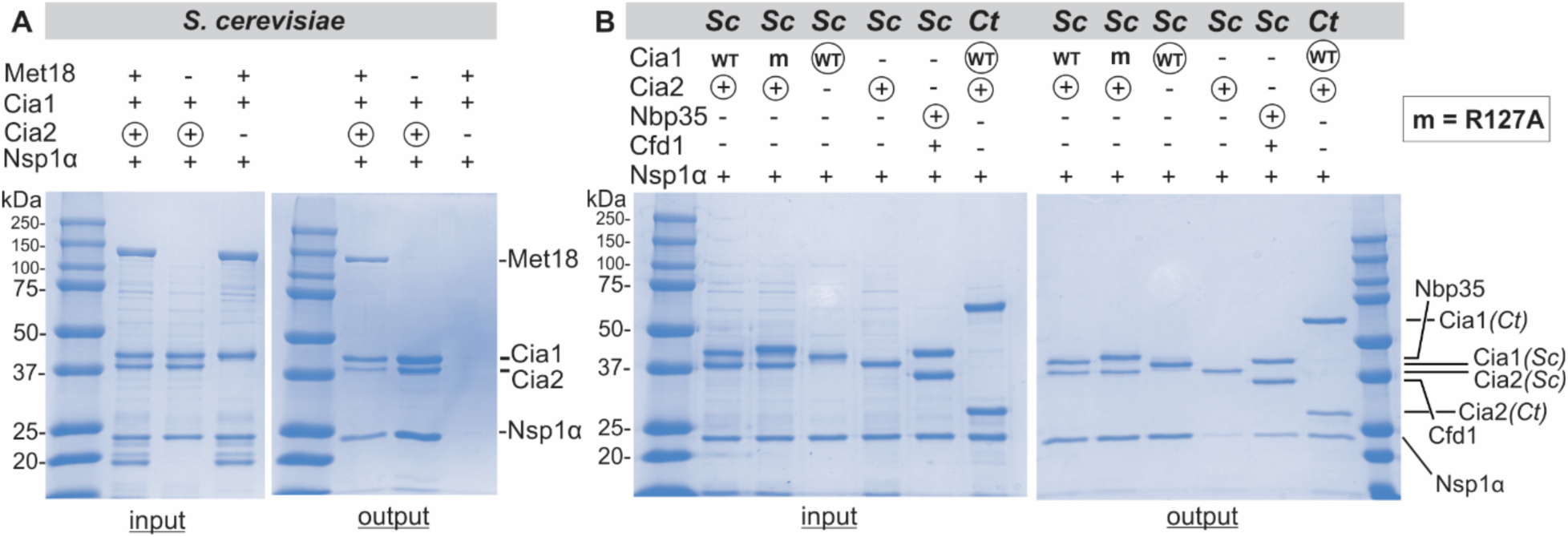
Nsp1α forms a complex with the components of the CTC machinery and Fe-S scaffold proteins. **A** SDS-PAGE of *in vitro* pull-down assay assessing Nsp1α interactions with components of the *Sc*CTC employing strep-tactin chromatography. Circles indicate the immobilized ‘bait’ protein that carries a Strep-Tag(II). **A** SDS-PAGE of in-vitro pull down assay assessing Nsp1a interactions with the *Sc* scaffold proteins Cfd1 and Nbp35 as well as the Cia1 R127A mutant, and *Ct* Cia1 and Cia2 proteins.

Next, we sought to define the binding determinants of the CTC-Nsp1α interaction, as endogenous CIA clients exhibit distinct dependencies on recognition sites distributed across the surface of the CTC (*37, 55, 56*). Using recombinant CIA factors, we found that Nsp1α coeluted with *Sc*Cia1 both in the presence and absence of *Sc*Cia2 (**Fig. 4B**). Because a substantial subset of endogenous CIA clients recruits the CTC via a conserved targeting complex recognition (TCR) motif (*57*), we next asked whether Nsp1α engages the same TCR peptide binding pocket at the Cia1-Cia2 interface (*35*). Consistent with its lack of a TCR motif, Nsp1α bound normally to the *Sc* R127A*-*Cia1 variant, which disrupts TCR peptide-mediated client binding. The viral protein also failed to displace a fluorescent TCR peptide probe from the *Chaetomium thermophilum* (*Ct*) Cia1-Cia2 complex, even though this ortholog was able to interact with Nsp1α (**Fig. 4B, Fig. S13**). Together, these data indicate that Nsp1α engages a conserved surface on Cia1 that is distinct from the TCR peptide binding pocket at the Cia1-Cia2 interface.

### Mammalian Cia1 recognizes Nsp1α via the acidic patch along its third beta propeller domain

To determine whether the interaction between Nsp1α and the CTC is conserved in mammalian systems, we examined the binding of Nsp1α to the human Cia1 (*Hs*Cia1) ortholog, CIAO1. Like the yeast and fungal orthologs, CIAO1 pulled down Nsp1α, demonstrating that the viral client is also recognized by the human CIA system (**Fig. 5A**). The conservation across fungal and mammalian orthologs suggests that Nsp1α engages an important and highly conserved surface of Cia1 (**Fig. 5B**).

**Figure 5.**
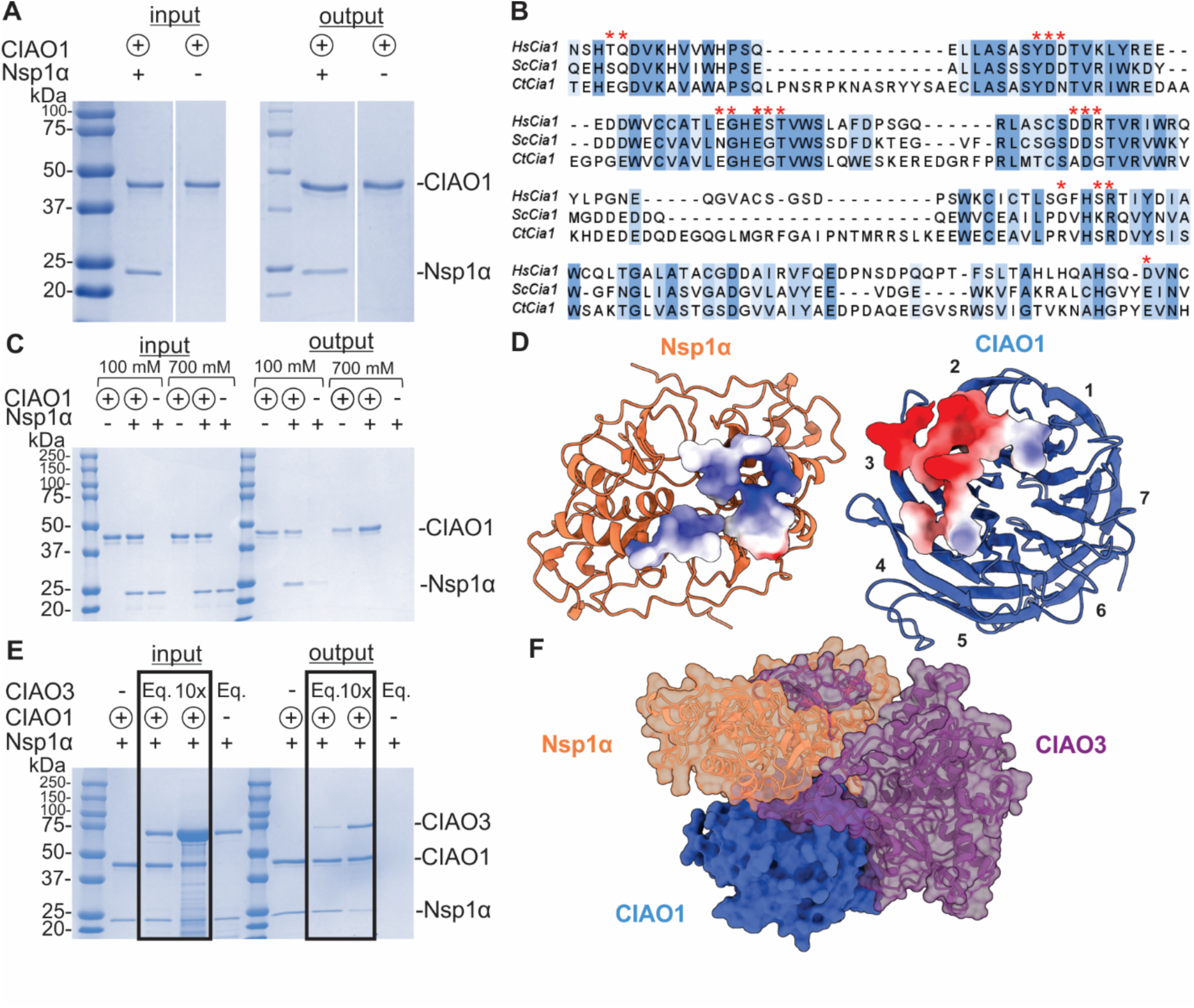
Nsp1α binding with the CTC is conserved and is mediated by electrostatic interactions. **A** *In vitro* pull-down of Nsp1α with strep-tagged CIAO1. **B** Multiple sequence alignment of *Hs*Cia1 (CIAO1), *Sc*Cia1, and *Ct*Cia1. Conserved residues are highlighted in blue, residues contacting Nsp1α in AlphaFold model within 3.5 Å are marked with a red asterisk. **C** *In vitro* pull-down of Nsp1α with CIAO1 under high ionic strength. **D** Electrostatic potential map of the binding interface on Nsp1α and CIAO1 (AlphaFold). **E** Competitive *in vitro* pull-down between Nsp1α and CIAO3 with CIAO1. **F** Superimposed AlphaFold model of CIAO1 complex with Nsp1α or CIAO3.

To gain molecular insights into how Nsp1α engages CIAO1, we examined the salt sensitivity of the CIAO1-Nsp1α interaction. Increasing the ionic strength from 100 to 700 mM abolished Nsp1α binding (**Fig. 5C**), indicating that complex formation is largely driven by electrostatic interactions. To define the structural basis of this interaction, we generated AlphaFold models of the CIAO1-Nsp1α complex, in which Nsp1α is a dimer as demonstrated in our experiments and consistent with previous reports (**Fig. S14, Fig. S15**) (*10, 18, 58*). These models position Nsp1α adjacent to an acidic patch on the highly conserved face of the third β-propeller of CIAO1 (**Fig. 5D**). Consistent with this arrangement, the surface of Nsp1α is enriched in basic residues, supporting an electrostatic mode of recognition.

We next examined whether Nsp1α engages CIAO1 in a manner similar to endogenous CIA client proteins (*37, 53*). CIAO3 has been proposed to act as an Fe-S cluster carrier that delivers nascent clusters to the CIA targeting complex and competes with client proteins for CIAO1 binding (*35, 36, 59*). Consistent with this model, increasing concentrations of CIAO3 led to a progressive reduction in the Nsp1α co-elution with human CIAO1 (**Fig. 5E**), indicating that these two proteins compete for overlapping binding sites. These observations suggest that Nsp1α interacts with CIAO1 through a conserved interface also utilized by endogenous CIA clients and the Fe-S carrier CIAO3, which is further supported by the AlphaFold model (**Fig. 5F**). Collectively, the data support the possibility that Nsp1α may access the CIA targeting complex via mechanisms analogous to host Fe-S proteins, with the potential to influence Fe-S cluster trafficking under conditions where Nsp1α is present at sufficient levels.

## Discussion

Nsp1α is the leader protease of PRRSV and a main component for sgRNA synthesis and host immune suppression (*8, 18*). Although it has been predicted to be a metalloprotein, the exact role of metal ions in its molecular function remains largely unexplored despite the fact that the putative ZF domain is central to its function. Disruption of the ZF motif is known to abolish sgRNA synthesis and immune suppression underscoring its functional significance (*11, 17–19*). These results highlight a distinct and essential regulatory role for the ZF domain in modulating host interactions and viral gene expression, marking it as an attractive therapeutic target.

We demonstrate that Nsp1α binds an Fe-S cluster at this ZF motif, redefining the metalloprotein identity of this key PRRSV enzyme. In agreement with previous studies, we found that mutations in any of the N-terminal ZF cysteines do not compromise proteolytic activity, protein stability, or dimerization (*10, 18*). Mössbauer, UV/VIS, and EPR spectroscopy identified the metallocluster as a [4Fe-4S]^2+^ cluster, challenging previous crystallographic data suggesting Zn^2+^ binding and providing a revised model for the metal-binding potential of Nsp1α. Our findings expand our understanding of the protein’s functionality and add to the growing evidence of the importance of Fe-S clusters in virus replication and pathogenicity (*24, 26, 38*).

Native Nsp1α, derived from expression of the parent Nsp1, does not strongly bind an Fe-S cluster (or Fe) irrespective of whether the protein is isolated aerobically or anaerobically. Nevertheless, Nsp1α purified from inclusion bodies is enriched in Fe-S clusters, suggesting that the metallocofactor is particularly O_2_-sensitive and destabilized post autoproteolytic cleavage. Semi-enzymatic reconstitution using IscS restored an intact [4Fe-4S]^2+^ cofactor in Nsp1α, but not in the ΔZF variant, confirming the N-terminal cysteines as ligands. Chemoproteomics further confirmed the binding site of the [4Fe-4S] ^2+^ as the N-terminal ZF motif. The [4Fe-4S]^2+^ cluster is highly labile under aerobic conditions and cannot be readily redox-cycled, only converting to a [3Fe–4S]^1^⁺ species upon oxidation. It should be noted that the [4Fe-4S] ^2+^ cluster may be reducible in the presence of an appropriate protein partner such as one of the many known host and viral interactors of Nsp1α (*60*). Although the precise redox potential remains unresolved, these data suggest that the cluster in Nsp1α belongs to the low-potential class (*44, 45*). Collectively, these characteristics resemble regulatory Fe-S centers such as those found in IRP1, NsrR and FNR suggesting a structural or sensing role for the Fe-S cluster in Nsp1α rather than in electron transfer (*61–64*).

To assess any functional consequences of metal binding, we examined how different cofactors affect the protease activity of Nsp1α. Expression of the parent Nsp1 under different metalation conditions had minimal impact on the extent of its cleavage *in cellulo*, indicating that this activity is largely metal-independent. This conclusion was reinforced by a variant that lacks the ZF cysteines, which retained protease activity. In contrast, mutation of either the C76 nucleophile or the H146 base, abolished cleavage and resulted in accumulation of insoluble, uncleaved Nsp1, confirming that proteolysis is intrinsic to Nsp1α and occurs *in cis*.

We further tested whether Nsp1α could recognize its substrate *in trans* either employing the catalytically dead Nsp1^C76A^ or a fluorogenic peptide encoding the cleavage site. While no cleavage of the Nsp1^C76A^ was observed, there was detectable, albeit slow, cleavage of the fluorogenic peptide. These results suggest that Nsp1α proteolytic activity is largely lost post-cleavage, and such domain specialization reflects a broader theme in viral non-structural proteins, including the PRRSV PLP2 and the pestivirus Ns2, which adopt multiple host-modulating roles following autoproteolytic processing (*49, 65*).

Despite its low basal protease activity, we leveraged it to probe the influence of the Zn and Fe-S metallocofactor. Zn had no effect at stoichiometric levels but inhibited protease activity in excess, in line with known Zn-mediated inhibition of cysteine proteases (*20, 21*). In contrast, [4Fe-4S] cluster binding enhanced protease activity four-fold relative to apo and Zn-bound forms. This enhancement although modest suggests a previously unrecognized regulatory function specific to the Fe-S cofactor in modulating Nsp1α activity.

The identification of Fe-S clusters in viral proteins is intriguing given that viruses lack the machinery to assemble these cofactors, implying a dependence on host pathways for their acquisition (*24, 25, 57*). *In vitro* pull-down experiments show that Nsp1α interacts with the early-stage cytosolic Fe-S assembly scaffold Nbp35-Cfd1 (*36*). These results indicate that Nsp1α engages components of the host Fe-S biogenesis machinery at an early step, providing a framework to understand its subsequent association with the downstream CTC. We also found that Nsp1α has a strong association with the yeast CTC comprising Cia1, Cia2, and Met18, with Cia1 emerging as the primary mediator of this interaction (*35, 36, 57*). This direct engagement with the host Fe-S cluster assembly system situates Nsp1α as a *bona fide* Fe-S protein, challenging earlier claims of Zn dependence. Comparable approaches have verified Fe-S clusters in other viral proteins, reinforcing their broader relevance in viral pathogenesis (*27–29*).

Biochemical assays and AlphaFold modeling support a model in which Nsp1α binds to Cia1 at the highly conserved patch on blade 3, an interface used by other Fe-S client proteins such as DNA2 helicase and DNA primase (*53*). In addition, this interaction is retained with both the human (CIAO1) and fungal (*Ct*Cia1) orthologs. Binding of Nsp1α to CIAO1 is mediated by electrostatic interactions and is disrupted under high-salt conditions. Nsp1α also competes with CIAO3 for this same site, suggesting that it may sequester CIAO1 from endogenous clients and perturb Fe-S cluster trafficking during infection (*36, 59*). This raises intriguing questions about how PRRSV, and Nsp1α in particular, may manipulate the host Fe-S cluster metabolism and whether this interference contributes to immune evasion or viral replication.

Altogether, our findings redefine Nsp1α as an Fe-S cluster protein that interacts with the host Fe-S assembly machinery. Given the strong interaction of Nsp1α with components of the CTC, we propose that Zn binding is likely an artifact of overexpression in heterologous systems rather than physiologically relevant, supporting the importance of reevaluating prior structural interpretations. Fe-S clusters, unlike Zn, are often multifunctional, influencing structure, catalysis, and redox behavior (*22, 23, 33, 64, 66*). This is evident in CPSF30, where Fe-S binding enhances RNA affinity, and in SARS-CoV-2 nsps, in which the Fe-S form enhances activity (*27–29, 67*). Similarly, the Fe-S cluster in Nsp1α may contribute to functional modulation, potentially regulating host interactions or immune evasion. The identification of an Fe-S cluster in Nsp1α not only redefines the molecular functions of this critical viral protein but also highlights an underappreciated vulnerability that may be exploited for the development of targeted antiviral therapies against PRRSV.

## Materials and Methods

### Materials

All chemicals were obtained from Fisher Scientific (unless specified otherwise) and were of high purity grade.

### Expression of Nsp1 and Nsp1α

Full length Nsp1^WT^ (aa: 1-382), Nsp1^ΔZF^ (C8,10,25,28A) (1–382), Nsp1^C70A^ (aa:1-382), Nsp1^C76A^ (aa:1-382), MBP-Nsp1^C76A^ (aa:1-382), Nsp1^C76A^-Nsp2 (aa:1-501), Nsp1^H146A^ (aa:1-382) or Nsp1α^WT^ (aa:1-180) carrying an N-terminal His-tag was inserted into a pET-28a+ vector which carries a kanamycin resistance, were all generated by Genscript USA Inc. (Piscataway, NJ). The Nsp1 sequence (NCBI accession code for Nsp1: NP_740595.1) was from the Porcine Reproductive and Respiratory syndrome Virus (PRRSV). The Nsp1/Nsp1α plasmids were transformed into Rosetta (DE3) T7 Express *Escherichia* (*E.*) *coli* competent cells (New England Biolabs, Ipswitch, MA). Cells were grown in minimal (M9) media containing 50 μg/ml kanamycin and either 125 μM ^56^Fe(NH_4_)_2_SO_4_ or no metals at 37 °C with shaking (200 rpm) until an OD_600_ ∼ 0.6 was reached, at which point they were transferred to 4 °C for 1 hr.

Protein expression was induced by addition of 0.5 mM isopropyl β-d-1-thiogalactopyranoside (IPTG). For Fe enrichment, cultures were supplemented with an additional 125 μM natural abundance ^56^Fe(NH₄)₂SO₄. For isotopic labeling, 125 μM ^57^Fe was added in place of ^56^Fe. For Zn enrichment, cultures were supplemented with 20 μM ZnSO₄. Following induction and supplementation, cells were incubated at 18 °C for 18-20 hr and harvested by centrifugation at 1,600 × g for 25 min at 4 °C. Cell pellets were flash-frozen in liquid nitrogen and stored at -80 °C until further use.

### Purification of native Nsp1α WT and variant apo proteins

Cell pellets were resuspended in lysis buffer (20 mM HEPES, 500 mM NaCl, and 20 mM imidazole, pH 7.5), and lysed by sonication at 4 °C. The lysate was centrifuged at 40,000 × *g* for 20 min to remove insoluble cell debris. For SDS-PAGE and western blot analysis, 20 μL of lysates were mixed 1:1 with 2× reducing sample buffer and heated at 95 °C for 5 min. Samples were resolved on 15% SDS–PAGE gels at 180 V for 55 min. Proteins were transferred to PVDF membranes using wet transfer (100 V, 60 min). Membranes were blocked in 5% non-fat milk in TBST for 1 hr and probed with an HRP-conjugated anti-His antibody (1:5,000). Bands were visualized by chemiluminescence.

The clarified lysate was applied to a Ni^2+^ chelating column (10 mL Ni^2+^-NTA agarose), followed by a 5 column-volume (CV) wash with lysis buffer. The bound protein was eluted with an eluting buffer (50 mM HEPES, 500 mM NaCl, 0.5 mM tris(2-carboxyethyl)phosphine (TCEP), and 250 mM imidazole, pH 8), then concentrated with a 30 kDa MWCO Amicon Centrifugal Filter (Millipore, Sigma-Aldrich). The concentrated protein was loaded onto a size exclusion S200 column (GE Healthcare), which was equilibrated with Storage Buffer (50 mM HEPES, 500 mM NaCl, 10% glycerol, 0.5 mM TCEP, pH 8.0). Nsp1α^apo^ was prepared by incubating the concentrated protein with 10 mM EDTA for 1 hr, prior to loading onto size exclusion column. Fractions containing pure protein were pooled and further concentrated, then flash frozen in liquid nitrogen and stored at -80 °C.

### Nsp1 Solubilization and Reconstitution

Five grams of *E. coli* cells expressing either Nsp1α^WT^, Nsp1^WT^ or the Nsp1^ΔZF^ variant were resuspended in 25 mL lysis buffer (as described in the native-purification section) inside an anaerobic chamber and lysed by sonication. Lysates were clarified by ultracentrifugation at 40,000 × g for 25 min. The insoluble pellets were resolubilized in 8 M urea, 20 mM CHES, 10 mM β-mercaptoethanol, and 1 mM EDTA (pH 9.0) using a Dounce homogenizer, then clarified by ultracentrifugation. Supernatants were applied to 1 mL Ni-NTA columns, washed with 10 CV of solubilization buffer, and eluted with the same buffer without EDTA and containing 250 mM imidazole. For reconstitution experiments, 1 mM DTT 125 µM ^57^Fe and Na₂S were added to purified protein and incubated for 1 hr before removal of unbound Fe and S on a Ni-NTA column with 10 CV wash of the solubilization buffer. In all cases, imidazole was removed by successive dilution and concentration in 10 kDa MWCO centrifugal concentrators into solubilization buffer without EDTA. All steps were performed at room temperature unless noted otherwise, with incubations at 4 °C in chilled buffers. Samples were frozen in Mössbauer cups by liquid nitrogen immersion.

### Expression and isolation of soluble Nsp1^C76A^

The variant Nsp1^C76A^ was inserted into the pMtac vector (kindly gifted by Dr. Michael Marr, Brandeis University, MA) which allows for its expression as a fusion with the maltose-binding protein (MBP) with a cleavable TEV recognition site (*68*). MBP-Nsp1^C76A^ was purified with 4 MBPTrap columns in tandem (5 mL, Cytiva) following previously reported protocols: Lysis and Wash Buffer (50 mM HEPES, 500 mM NaCl, 0.5 mM TCEP, pH 8.0), Elution Buffer (50 mM HEPES, 500 mM NaCl, 10 mM Maltose, 0.5 mM TCEP, pH 8.0) (31). The eluted protein was concentrated with a 30 kDa MWCO Amicon Centrifugal Filter (Millipore, Sigma-Aldrich). The concentrated protein was loaded onto a size exclusion S200 column (GE Healthcare), which was equilibrated with Storage Buffer (50 mM HEPES, 500 mM NaCl, 10% glycerol, 0.5 mM TCEP, pH 8.0). The MBP tag was cleaved from the fusion protein using TEV protease at a 1:10 ratio (TEV:MBP-Nsp1^C76A^) overnight at 4 °C. The cleaved protein was confirmed by SDS-PAGE and loaded onto a size exclusion S200 column (GE Healthcare), which was equilibrated with Storage Buffer (50 mM HEPES, 500 mM NaCl, 10% glycerol, 0.5 mM TCEP, pH 8.0). The fractions containing pure Nsp1^C76A^ were pooled and further concentrated, then flash frozen in liquid nitrogen and stored at -80 °C.

### Expression of SARS-CoV PL^pro^ and PRRSV PLP2

SARS-CoV PL^pro^ (NP_828862) carrying an N-terminal His-tag was inserted into a pET-28a+ vector which carries a kanamycin resistance was generated by Genscript USA Inc. (Piscataway, NJ). The PLP2 domain from PRRSV DV was inserted in a pASK vector as previously described affording the pASK-DV_399-578_ plasmid (*49*). Expression and purification of the plasmids of PL^pro^ and PLP2 were performed as previously described.

### IscS-mediated Fe-S reconstitution

The *E. coli* IscS was expressed as a fusion with an N-terminal SUMO tag (the plasmid of pSUMO-IscS encodes for kanamycin resistance and was kindly gifted by Squire S. Booker) and isolated as described previously (*46*).Protein to be reconstituted was degassed using a vacuum Schlenk line sparged with Argon. The degassed protein (400 μM) was incubated with 5 mM Dithiothreitol (DTT) for 30 min. Fe(II) was added in the form of Fe(NH_4_)_2_SO_4_ was in four equal portions over the course of 1 hr to final concentration of 1.6 mM. Fe-S cluster reconstitution was initiated when L-cysteine was added to 5 mM final concentration, followed by the cysteine desulfurase IscS at 5 μM final concentration and left in the chamber overnight at 4 °C. The resulting solution was centrifuged at 20,000 x g and the supernatant passed twice over PD-10 desalting columns (Cytiva) to remove any non-bound iron and sulfide.

### Protease assays

All fluorescence measurements were carried out in a Biotek Cytation 5 plate reader (Agilent) in 96-well plates. For each condition, 50 µL of enzyme solution was dispensed in triplicate into a black 96-well microplate, followed by addition of 50 µL of substrate solution. Plates were preheated to 25 °C prior to initiating the assay. Measurements were performed at a setpoint of 25 °C with no applied gradient.

Protease activity of Nsp1α was assessed using a generic protease substrate FITC-casein. Readout was collected in endpoint mode using the following filter set: excitation 485 ± 20 nm, emission 535 ± 20 nm, top-read optics, and extended gain. The light source was a xenon flash lamp at high energy with extended dynamic range enabled. Enzyme samples were diluted to 30 µM in assay buffer (50 mM CHES, 500 mM NaCl, pH 9.0), and FITC-casein substrate was diluted to 1 µM in the same buffer immediately prior to use.

Deubiquitinase activity of Nsp1α was measured using the peptide substrate Z-RLRGG-AMC and Ubiquitin-AMC using the following filter site: excitation 340 ± 20 nm, emission 450 ± 20 nm, top-read optics, gain set to 50.

Activity of Nsp1α was also interrogated with a fluorogenic peptide incorporating the internal cleavage sequence of Nsp1α (FECAMATVYD) that was synthesized by Lifetein. The substrate incorporates the fluorophore 5-[(2-aminoethyl)amino]naphthalene-1-sulfonic acid (EDANS) at the N-terminal end of the sequence and an EDANS-quenching moiety (Dabcyl, (4-(4-dimethylaminophenylazo)-benzoic acid)) on the C-terminal end. Following cleavage, the fluorophore EDANS is released yielding fluorescence that can be detected using excitation wavelengths at 360 ± 20 nm and emission wavelengths at 500 ± 20 nm. Nsp1α^apo^, Nsp1α^Fe-S^, and Nsp1α^Zn^ were prepared as follows. Nsp1α^apo^ was prepared directly in assay buffer. For Nsp1α^Zn^, ZnCl₂ was added at one molar equivalent relative to protein concentration and incubated for 20 min. For Nsp1α^Fe-S^, assays were prepared inside an anaerobic chamber, and plates were immediately sealed with a clear adhesive film to minimize oxygen penetration during the measurements. Fluorescence was monitored over the course of 1 hr with 1 min intervals.

### Mössbauer Spectroscopy

Mössbauer spectra were recorded on a WEB Research spectrometer (Edina, MN) equipped with a Janis SVT-400 variable-temperature cryostat. Zero velocity was calibrated as the centroid of the spectrum of a-Fe recorded at room temperature. WMOSS spectral analysis software (www.wmoss.org, WEB Research, Edina, MN) was used to simulate and analyze Mössbauer spectra.

### Protein and Metal quantification

All protein concentrations were determined by the A280 and calculated extinction coefficient using ProtParam (expasy.org). Nsp1α^WT^ (24450 M^-1^·cm^-1^), Nsp1α^ΔZF^ (23950 M^-1^·cm^-^_1_), Nsp1^C76A^ (42587 M^-1^·cm^-1^), Nsp1^H146A^ (42521 M^-1^·cm^-1^), Nsp1α^C70A^ (23950 M^-1^·cm^-1^), MBP-Nsp1^C76A^ (128230 M^-1^·cm^-1^), Nsp1^C76A^-Nsp2 (78840 M^-1^·cm^-1^). Metal analysis of the as-purified proteins was determined by inductively coupled atomic emission spectrometry (ICP-AES, Laboratory for Isotopes and Metals, Pennsylvania State University).

### Isotopic NEM-labeling

100 µL of Nsp1α (50 µM) was incubated with either 1 mM of isotopically light NEM-d_0_ (Nsp1α^apo^) or heavy NEM-d_5_ (Nsp1α^Zn^ or Nsp1α^Fe-S^), mixed and incubated at room temperature for 1 hr. The protein samples were frozen and stored at -80 °C until further processing. A detailed sample preparation protocol can be found in the SI along with Mass-spectrometry and data processing.

### Phylogenetic Analyses

We selected 30 unique Nsp1α protein sequences that represent the bulk of the Arteriviruses in which Nsp1α occurs. The sequences were aligned with the MAFFT software(*69*) and the maximum-likelihood rooted phylogenetic tree was subsequently computed with the IQ-tree software (*70*)using the PMB+I+G4 substitution model, selected as the best-fit model on the basis of the Bayesian Information Criterion (BIC). Branch support was assessed using the SH-like approximate likelihood ratio test (aLRT) with 1000 replicates and standard bootstrap analysis. The resulting tree was visualized using Interactive Tree of Life (iTOL) and rooted with the Wobbly Possum Virus Nsp1α sequence as the outgroup.

### In-vitro pull-down

All experiments were performed at 4 °C. Bait-only and prey-only controls were performed in parallel as indicated (*35*). An affinity-tagged bait protein (Strep tag) was mixed with an equimolar amount of one or more prey proteins for 1 hr.

Strep-tagged bait proteins (10 μM final concentration for *Sc*Cia2; 8 μM for CIAO1, *Ct*Cia1-^Strep^Cia2, or *Sc* ^Strep^Nbp35-Cfd1) were mixed with equimolar concentrations of Nsp1α and incubated in assay Buffer [ 50 mM Tris, 100 mM NaCl, 5% glycerol, 5 mM BME, pH 8.0]. After incubation for 1 hr, the mixture (∼630 µL) was then added to Strep-Tactin XT Superflow (200 µL, IBA) resin and incubated for 1 hr with gentle rocking. The resin was collected, washed, and eluted with Assay Buffer supplemented with 50 mM D-Biotin and 100 mM NaOH. The resulting elution fractions were analyzed along with input samples via a 12% SDS-PAGE gel. Each copurification analysis included parallel controls: input samples to verify equal variant concentrations and either a “no-bait” or “non-CIA bait” (PPAR-γ) control to detect nonspecific prey-resin and prey-bait binding.

To determine the effect of buffer’s ionic strength on protein-protein interaction, proteins were incubated in the assay buffer where NaCl concentration was increased up to 700 mM [50 mM Tris, 700 mM NaCl, 5% glycerol, 5 mM BME, pH 8.0]. For the competitive affinity co-purification assay between Nsp1α, CIAO3, and CIAO1, besides the equimolar mixture, a reaction with a 10-fold molar excess of CIAO3 over CIAO1 was prepared. Elution and SDS-PAGE analysis of elution fractions was carried out as outlined above.

### Competitive fluorescence anisotropy (FA) assay

For the competitive FA assay, *Ct*Cia1-Cia2 complex (5 μM) was incubated with 0.05 μM of FITC-HLDW, followed by titration of Nsp1α (0-40 μM) in a NaPi-FA buffer (50 mM Na_2_HPO_4_ (pH 8.0), 100 mM NaCl, 5% glycerol, 5 mM β-mercaptoethanol). FA measurements were collected as previously described (*35*).

### Stopped-flow Absorption experiments (SF-Abs)

Stopped-flow experiments were performed with the Applied Photophysics SX-20 instrument (Leatherhead, UK) equipped with a temperature control unit set at 5 °C. The unit was housed in an anoxic chamber (CoyLab). In single-mix (two-syringe) experiments, a solution containing 100 μM of [4Fe-4S]^2+^ cluster enriched Nsp1α was reacted 1:1 with 1.8 mM O_2_-saturated buffer. The optical path length used was 10 mm. For the characterization of the reactions, the photodiode array detector (PDA) was used to acquire time-resolved absorption spectra.

### EPR Spectroscopy

All EPR samples were prepared in Storage Buffer under O_2_-free conditions in an anaerobic glovebox. Samples were reduced with 6 mM sodium dithionite for 20 min at room temperature prior to being frozen in liquid N_2_, unless stated otherwise.

X-band continuous-wave (CW) EPR measurements were performed on a Bruker EleXsys E500 EPR spectrometer (operating at approximately 9.38 GHz) equipped with a rectangular resonator (TE102) and a continuous-flow cryostat (Oxford 910) with a temperature controller (Oxford ITC 503). The CW EPR spectra were recorded at 10 K, with a microwave power of 0.2 mW and a modulation amplitude of 1 mT, unless stated otherwise.

### Isothermal titration calorimetry (ITC)

Protein samples were dialyzed against a buffer containing 50 mM HEPES, 500 mM NaCl, pH 8. ITC experiments were carried out in a Nano ITC instrument (TA Instruments) at 5 °C. The titrations were performed by injecting 1.5 μL aliquots of 1 mM ZnCl_2_ into the calorimeter cell containing a 350 μL solution of either Nsp1α^WT^ or Nsp1α^ΔZF^ (50 μM) with a constant stirring speed at 150 rpm. The data were analyzed with the NanoAnalyze software using the independent fit model. All the uncertainties were estimated by the built-in statistics module with 1000 synthetic trials and 95% confidence level.

## Acknowledgments

The authors thank Dr. Paul Ralifo for providing us access to the EPR spectrometer at the Chemical Instrumentation Center of Boston University (Boston, MA). The authors thank Dr. Laura J. Liermann for ICP-AES analysis at the Laboratory for Isotopes and Metals in the Environment at Pennsylvania State University (University Park, PA). The authors are also thankful to Alex MacNeil and Dr. Douglas L. Theobald for providing unrestricted access to the Cytation 5 Plate Reader. The authors thank Dr. Bruce Goode (Brandeis University) for providing unrestricted access to their microfluidizer.

## Competing Interest Statement

The authors declare no competing interests.

## Funding Sources

This work was supported by the National Institutes of Health (R01-GM126303 and R35-GM156452 to M.-E.P.; R01-GM121673 to D.L.P.; R35-GM134964 to E.W.). Trent Quist was supported by a predoctoral fellowship (T32 GM135126).

## Notes

### Competing Interest Statement

The authors have declared no competing interest.

