## Supporting Information for "Discovery of a [4Fe-4S] cluster in the PRRSV Nsp1α leader protease reveals host-virus interplay in its downstream functions"

**Mass Spectrometry Sample preparation**

Frozen, labeled protein samples were thawed on ice and immediately precipitated with 5 % TCA (5 µL of 100 % TCA in water). After addition of TCA, samples were vortexed and stored at -80 ºC for at least 2 hr or overnight. Samples were thawed on ice, then centrifuged at 15,000 g for 10 min at 4 ºC to collect precipitated protein. The supernatant was removed, and the protein pellet was resuspended in 500 µL of ice-cold acetone by vortexing and sonication. The protein precipitate was pelleted by centrifugation at 5,000 g for 10 min at 4 ºC. The supernatant was removed, and the protein pellet was allowed to air-dry until all acetone was evaporated. The protein precipitate was resolubilized in 30 µL of urea (fresh 8 M stock in PBS) by sonication, followed by the addition of 70 µL of ammonium bicarbonate (100 mM fresh stock in water). To reduce protein thiols, 1.5 µL of DTT (150 mg/mL fresh stock in water) was added and samples were then incubated at 75 ºC for 15 min. To alkylate free thiols, 2.5 µL of iodoacetamide (94 mg/mL fresh stock in water) was added to samples, followed by incubation at room temperature for 30 min in the dark. The protein samples were then diluted with 120 µL of PBS. To digest protein samples, 2.5 µL of calcium chloride (100 mM stock in water) and 4 µL of sequencing grade trypsin (resuspended at 20 μM in trypsin resuspension buffer) were added and the samples were incubated overnight at 37 ºC. The samples were acidified by the addition of 12.5 µL of MS-grade formic acid and combined pairwise (light NEM-d_0_ labeled Nsp1α^apo^ with heavy NEM-d_5_ labeled Nsp1α^Fe-S^ or Nsp1α^Zn^) into a clean eppendorf tube (final volume ~ 500 µL). Samples were desalted using Sep-Pak C18 cartridges, which had been conditioned with 3 x 1 mL 100 % MeCN, then equilibrated with 3 x 1 mL Buffer A (5 % MeCN, 0.1 % formic acid). Samples were loaded onto the cartridges and allowed to flow through under ambient pressure. Samples were then reloaded onto the cartridge to improve yield. Cartridges were washed with 3 x 1 mL Buffer A. Samples were eluted from the cartridge into LoBind eppendorf tubes using 2 x 0.5 mL Buffer B (80 % MeCN, 0.1 % formic acid). Samples were evaporated to dryness in a SpeedVac and stored at -20 ºC until LC-MS/MS analysis. Samples were resuspended in 1 mL of Buffer A.

**LC-MS/MS Analysis**.

Mass spectrometry-based proteomic analysis of three biological replicates was performed in technical duplicate. Samples were analyzed by LC-MS/MS on an Orbitrap Exploris 240 mass spectrometer running Xcalibur v4.4 (Thermo Scientific) coupled to a Dionex Ultimate 3000 RSLCnano system.. Samples (5 µL) were injected onto an Acclaim PepMap 100 (Thermo 164750) loading column. Peptides were eluted onto an Acclaim PepMap RSLC (Thermo 164534) and separated with a 1 hr gradient of Buffer Buffer B (80 % MeCN, 0.1 % formic acid) in Buffer C (100 % H_2_O, 0.1 % formic acid) at a flow rate of 0.3 µL/min. The spray voltage was set to 2.1 kV. One full MS1 scan (120,000 resolution, 350-1800 m/z, RF lens 65 %, AGC target 300 %, automatic maximum injection time, profile mode) was obtained every 2 s with dynamic exclusion (repeat count 2, duration 10 s), isotopic exclusion (assigned), and apex detection (30 % desired apex window) enabled. A variable number of MS2 scans (15,000 resolution, AGC 75 %, maximum injection time 100 ms, centroid mode) were obtained between each MS1 scan based on the highest precursor masses, filtered for monoisotopic peak determination, theoretical precursor isotopic envelope fit, intensity (5 x 10^4^), and charge state (2-6). MS2 analysis consisted of the isolation of precursor ions (isolation window 2 m/z) followed by higher-energy collision dissociation (HCD) (collision energy 30%).

**Mass Spectrometry Data Processing.**

RAW files were uploaded to the Thermo Proteome Discoverer v2.4 software package and searched using the SequestHT algorithm against a UniprotKB database containing both the *E. coli* K12 proteome and the WT Nsp1α protein sequence. Trypsin was specified as the protease with a maximum of two missed cleavages. Peptide precursor mass tolerance was set to 10 ppm with a fragment mass tolerance of 0.02 Da. Oxidation of methionine and modification of cysteine by isotopically light (+125.048) or heavy (+130.079) NEM were set as dynamic modifications, while acetylation and/or methionine-loss of the protein N-terminus were set as static modifications. The false discovery rate (FDR) for peptide identification was set to 1 %. Light/heavy (L/H) ratios for cysteine containing Nsp1α peptides were calculated as the ratio of the NEM-d_0_ over NEM-d_5_ modified peptide precursor ion intensities.

**Table S1** Protein sequences used in Nsp1α multiple sequence alignment and in the generation of the phylogenetic tree.

| **NCBI Accession ID** | **Organism** |
| --- | --- |
| YP_009505547.1 | Porcine reproductive and respiratory syndrome virus 2 |
| NP_047406.1 | Porcine reproductive and respiratory syndrome virus |
| YP_009337023.1 | Rat arterivirus 1 |
| NP_042572.1 | Lactate dehydrogenase-elevating virus |
| YP_009118961.1 | African pouched rat arterivirus |
| YP_009388591.1 | Olivier's shrew virus 1 |
| YP_009067051.1 | Mikumi yellow baboon virus 1 |
| YP_009221994.1 | Kafue kinda chacma baboon virus |
| NP_203542.2 | Simian hemorrhagic fever virus |
| YP_009140476.1 | Pebjah virus |
| YP_009121773.1 | DeBrazza's monkey arterivirus |
| YP_009505556.1 | Kibale red-tailed guenon virus 1 |
| YP_009249808.1 | Free State vervet virus |
| YP_009505568.1 | Simian hemorrhagic encephalitis virus |
| YP_009824945.1 | Zambian malbrouck virus 1 |
| YP_009344805.1 | Kibale red colobus virus 1 |
| YP_009130632.2 | Wobbly possum disease virus |
| NP_127507.1 | Equine arteritis virus |
| QAB05846.1 | South west baboon virus |
| YP_009362004.1 | Kibale red colobus 2 virus |
| WPV62898.**1** | Wufeng rodent 1 virus |
| WPV62906.1 | Wufeng rodent 2 virus |
| WPV62879.**1** | Jingmen shrew 1 virus |
| QYL35101.1 | Praja virus |
| XNX57408.1 | Hedgehog arterivirus |
| WLW38111.1 | Oecomys arterivirus |
| ATP66644.1 | RtClan arterivirus |
| WFD49970.1 | Bamboo rat arterivirus |
| QYL35081.1 | Lopma virus |
| APT40620.1 | RtClon arterivirus |

**Table S2** Metal content of Nsp1α purified from M9 media supplemented with different divalent metals as determined by ICP-AES.

| **Growth media** | **Fe molar eq** | **Zn molar eq** |
| --- | --- | --- |
| M9 no metals added | 0.00 | 0.09 |
| M9 + Fe(NH_4_)_2_SO_4_ | 0.00 | 0.50 |
| M9 + ZnSO_4_ | 0.02 | 1.00 |
| M9 + ZnSO_4_ & Fe(NH_4_)_2_SO_4_ | 0.02 | 0.49 |

**Table S3** Quantification of cysteine-containing Nsp1α peptides labeled with N-ethylmaleimide (NEM) under apo and 4Fe-4S conditions. Reported are L/H ratios across six replicates with mean and standard deviation.

| ***Nsp1α Peptide Sequence*** | ***Peptide Modifications*** | ***Cys*** | ***Nsp1α L/H Ratio (Apo / Zn)*** | | | | | | | |
| --- | --- | --- | --- | --- | --- | --- | --- | --- | --- | --- |
|  |  |  | ***RRep 1*** | ***RRep 2*** | ***RRep 3*** | ***RRep 4*** | ***RRep 5*** | ***RRep 6*** | **AAvg** | **ss.d.** |
| [R].CTCTPNAR.[V] | 2xNEM  [C1; C3] | 8,10 | 7.80 | 7.51 | 5.75 | 4.90 | 3.89 | 3.62 | **5.57** | 1.780 |
| [R].VFVAEGQVYCTR.[C] | 1xNEM  [C10] | 25 | 7.37 | 7.24 | 2.90 | 4.61 | 5.40 | 4.14 | **5.28** | 1.768 |
| [R].AFPTVECSPAGACWLSAIFPIAR.[M] | 2xNEM  [C7; C13] | 70,76 | 1.49 | 1.53 | 1.51 | 1.20 | 1.31 | 1.41 | **1.41** | 0.130 |

**Table S4** Quantification of cysteine-containing Nsp1α peptides labeled with N-ethylmaleimide (NEM) under apo and Zn conditions. Reported are L/H ratios across six replicates with mean and standard deviation.

| ***Nsp1α Peptide Sequence*** | ***Peptide Modifications*** | ***Cys*** | ***Nsp1α L/H Ratio (Apo / Zn)*** | | | | | | | |
| --- | --- | --- | --- | --- | --- | --- | --- | --- | --- | --- |
|  |  |  | ***RRep 1*** | ***RRep 2*** | ***RRep 3*** | ***RRep 4*** | ***RRep 5*** | ***RRep 6*** | **AAvg** | **ss.d.** |
| [R].CTCTPNAR.[V] | 2xNEM [C1; C3] | 8, 10 | 3.16 | 3.14 | 4.71 | 4.32 | 3.46 | 3.75 | **3.76** | 0.6397 |
| [R].VFVAEGQVYCTR.[C] | 1xNEM [C10] | 25 | 5.76 | 6.72 | 5.85 | 4.80 | 5.57 | 4.82 | **5.59** | 0.7186 |
| [R].AFPTVECSPAGACWLSAIFPIAR.[M] | 2xNEM  [C7; C13] | 70, 76 | 6.40 | 6.20 | 8.93 | 9.00 | 7.33 | 8.06 | **7.65** | 1.2153 |

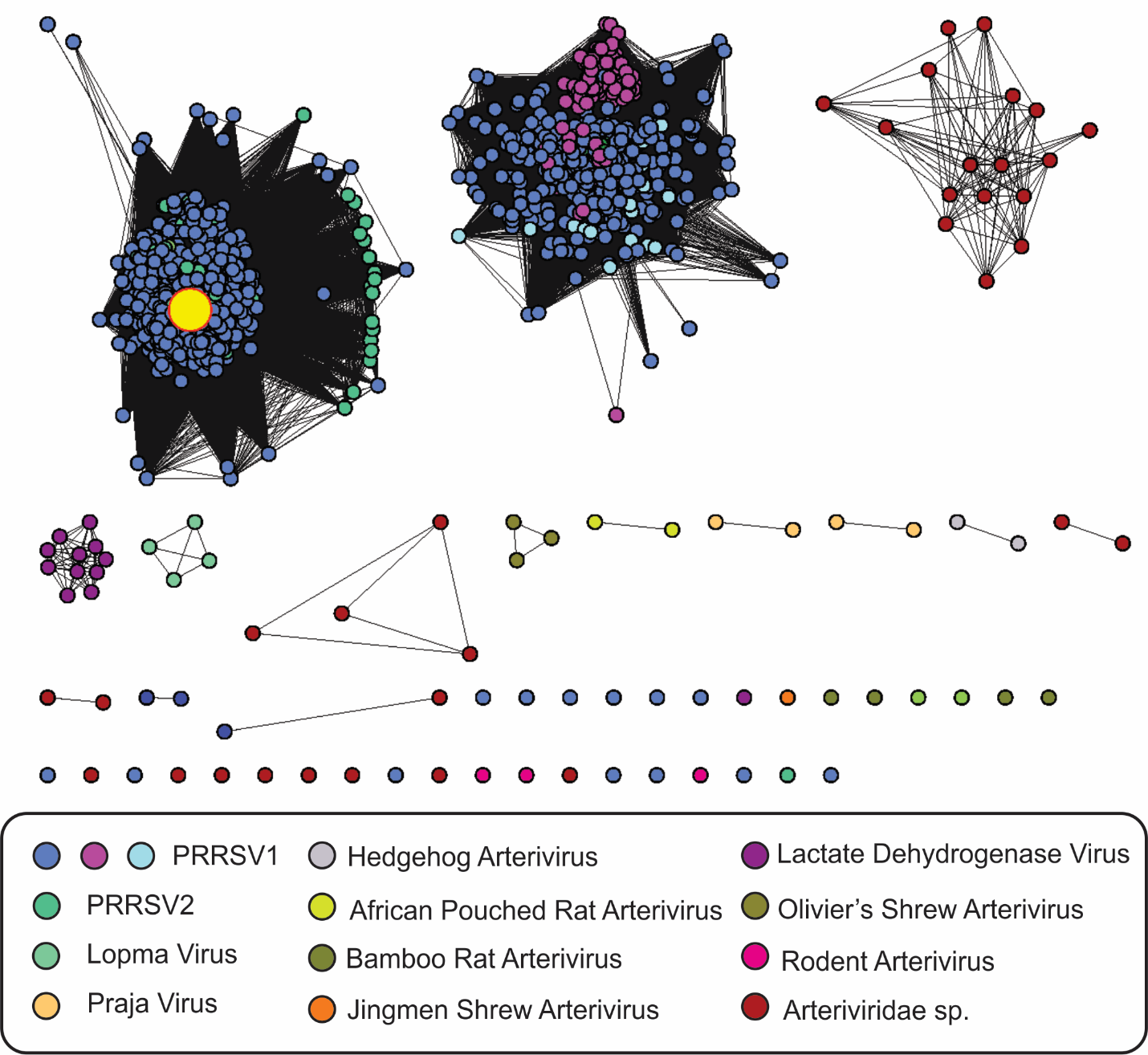

**Figure S1.** SSN of Nsp1α protein sequences retrieved from a blast search using the PRRSV1 Nsp1α sequence as an input. The SSN was generated using the web-based Enzyme Function Initiative-Enzyme Similarity Tool (EFI-EST) (1). The SSN was calculated with an alignment score of 95 and further refined to contain sequences between 170 and 180 amino acids long. The final SSN contains 2,231 unique sequences and the nodes have been colored according to the virus family. The large yellow circle indicates the input Nsp1α protein sequence.

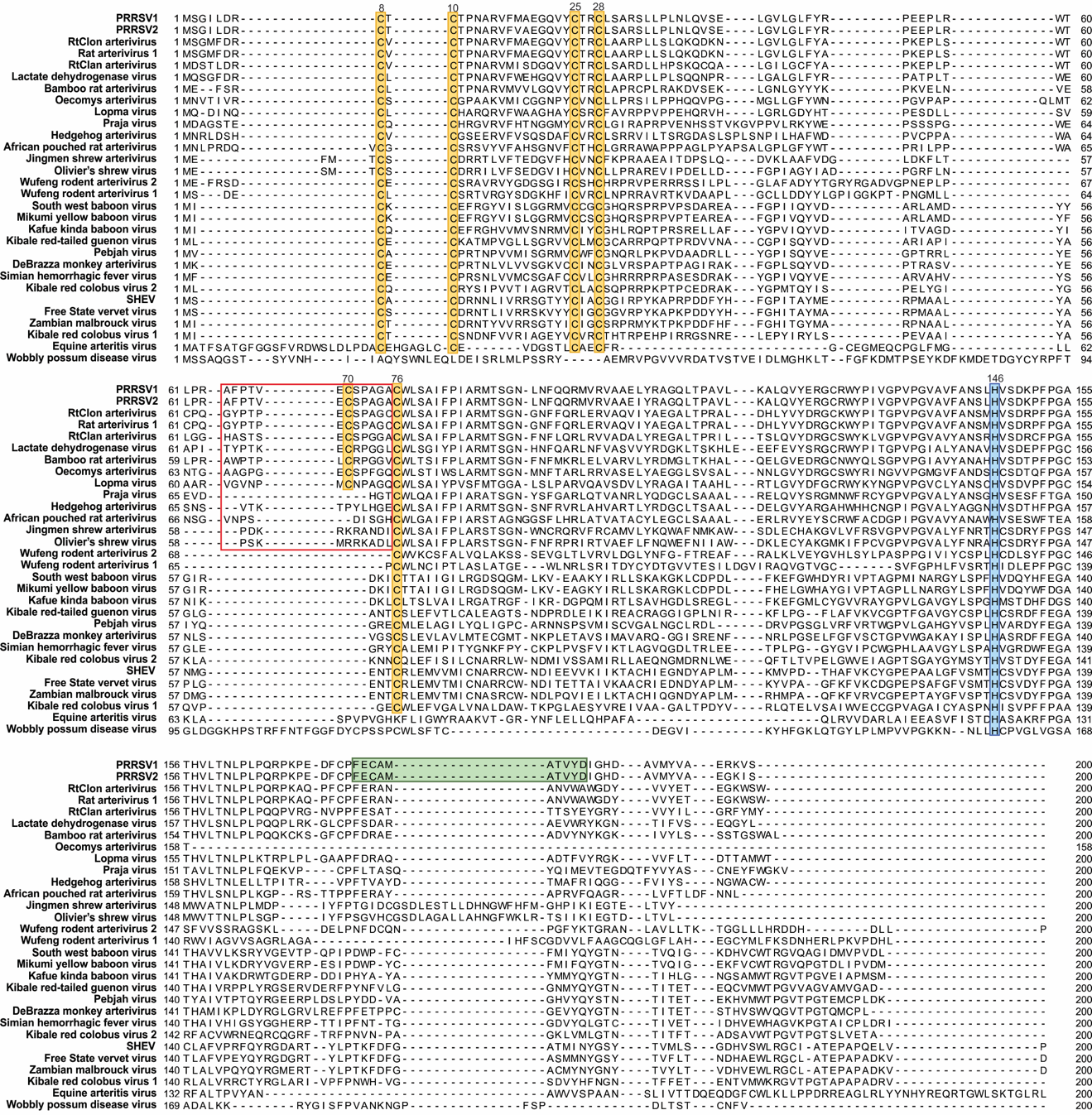

**Figure S2**. Multiple sequence alignment of Nsp1α protein sequences from phylogenetically diverse Arteriviruses. Conserved cysteine residues are highlighted in yellow, and histidine in blue. The loop insertion that contains the C70 metal binding residue is boxed in red and the internal cleavage sequence of PRRSV Nsp1α is shaded in green.

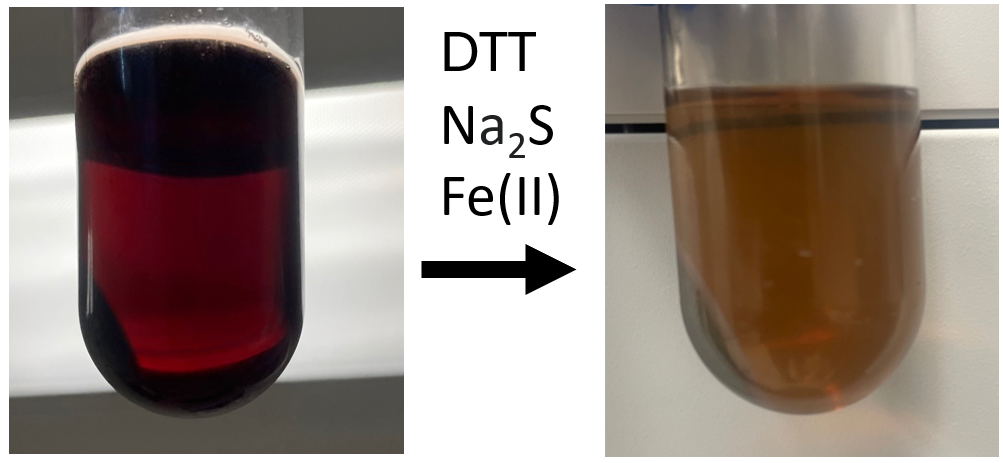

**Figure S3.** Wild type Nsp1α resolubilized from inclusion bodies in a buffer containing 8 M urea, 10 mM beta-mercaptoethanol, 20 mM CHES pH 9 before and after chemical reconstitution by the addition of 1mM DTT, 125 µM Fe(NH_4_)_2_SO_4_ and 125 µM sodium sulfide (Na_2_S) under O_2_-free conditions in an anaerobic glovebox.

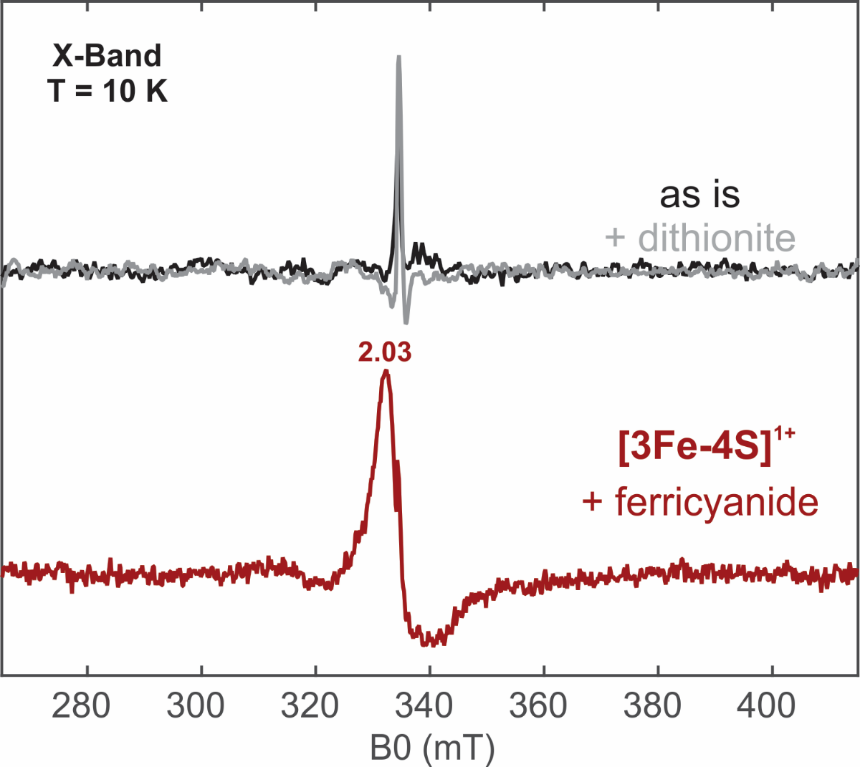

**Figure S4.** X-band continuous wave (CW) EPR spectra of chemically reconstituted Nsp1α^WT^ under different redox conditions. (Top) Nsp1α as-isolated (black trace) and treated with 10 mM sodium dithionite for 30 min (gray trace). (Bottom) Nsp1α treated with 10 mM potassium ferricyanide for 30 min. Experimental conditions: microwave frequency = 9.36 GHz, T = 10 K, microwave power = 0.64 mW, modulation amplitude = 1 mT.

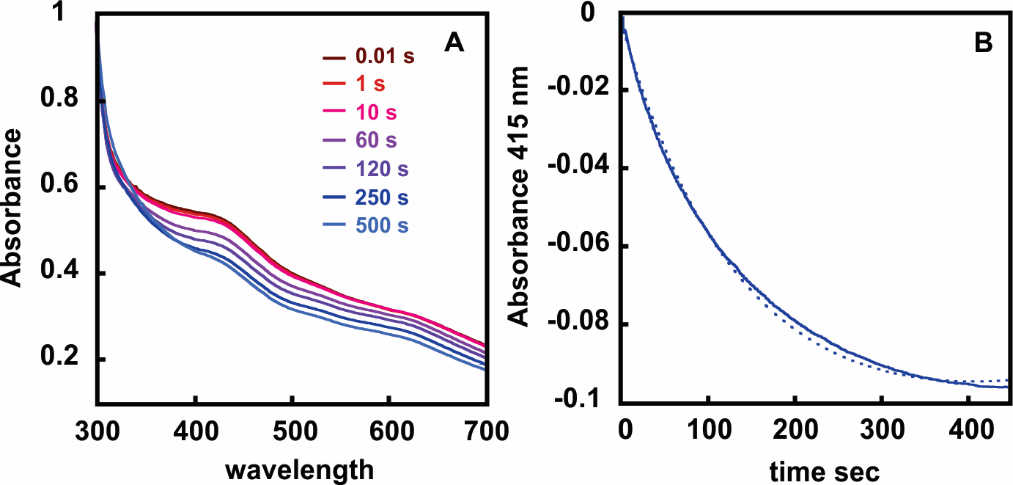

**Figure S5.** Stopped-flow absorption kinetic experiments monitoring the oxygen sensitivity of the Fe-S cluster in semi-enzymatically reconstituted Nsp1α^WT^. (A) Absorption spectra recorded after rapid mixing of Nsp1α^WT^ (0.1 mM) with an equal volume of O_2_-saturated buffer (1.8 mM) in a 1:1 ratio. (B) The kinetics of the decay of the [4Fe-4S] cluster signal signifying its decomposition were monitored by following the absorbance at 412 nm (solid trace). The fit to a single exponential decay has been included as a dashed line from which the rate of the decay was estimated. The experiments were carried out at 5 °C.

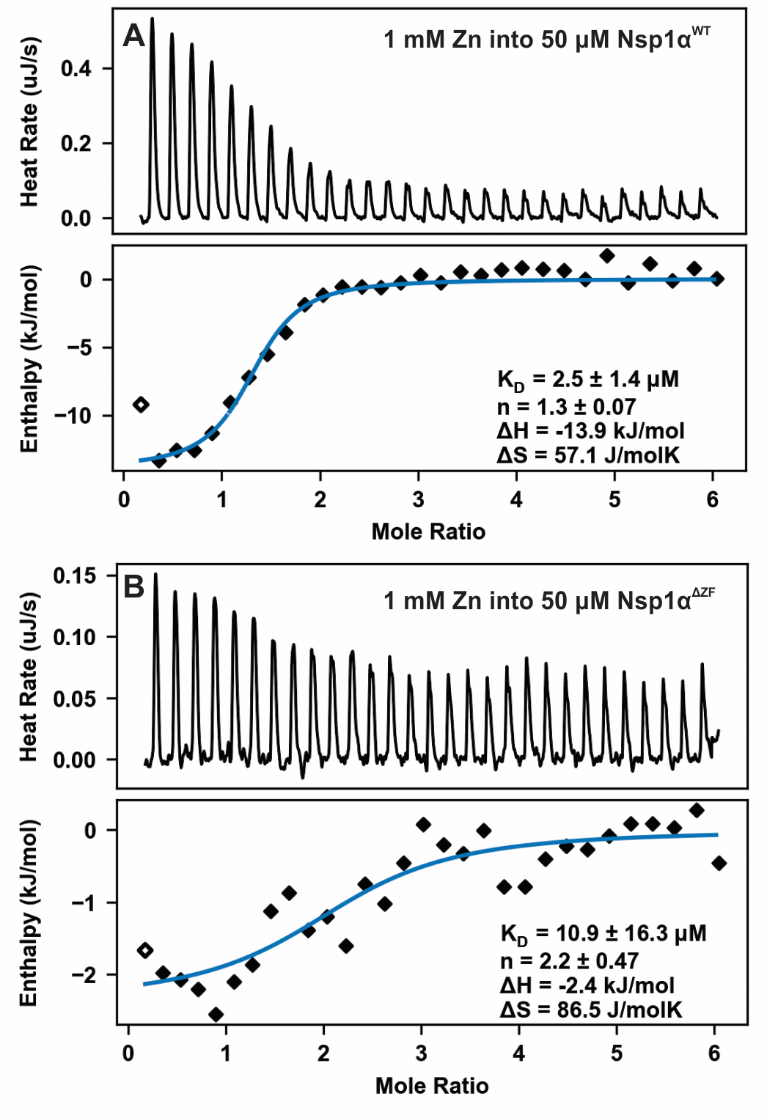

**Figure S6.** Isotherms for wild type Nsp1α (A) and Nsp1α^ΔZF^ (B) titrated with 1 mM ZnCl_2._ The data were analyzed with the NanoAnalyze software using the independent fit model. All the uncertainties were estimated by the native Statistics module with 1,000 synthetic trials and at 95 % confidence level.

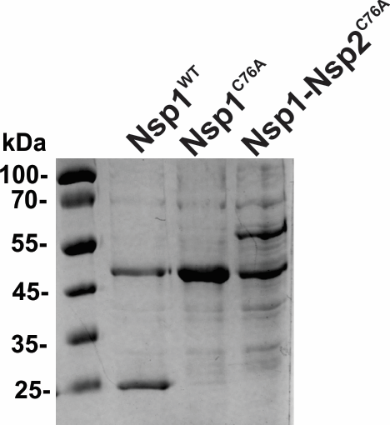

**Figure S7.** Auto-proteolytic activity of Nsp1 constructs in cell lysates. Cleavage products were resolved by SDS-PAGE and visualized by Coomassie staining. In the case of Nsp1^WT^ there are two bands, one corresponding to the uncleaved parent protein Nsp1 (46.4 kDa) and one to that of the Nsp1α/Nsp1β products (23.7/22.7 kDa respectively). Mutation of C76 to alanine abolishes proteolytic cleavage and only the band of the parent Nsp1^C76A^ protein is detectable. Expression of Nsp1^C76A^-Nsp2 (1:501 aa, mw = 59.6 kDa) in cell lysates results in the appearance of two bands, one corresponding to the parent protein and a second lower molecular weight one that corresponds to the Nsp1^C76A^ protein (46.4 kDa), demonstrating that release of Nsp1α does not inhibit the proteolytic activity of Nsp1β that cleaves itself from the Nsp1-Nsp2 junction.

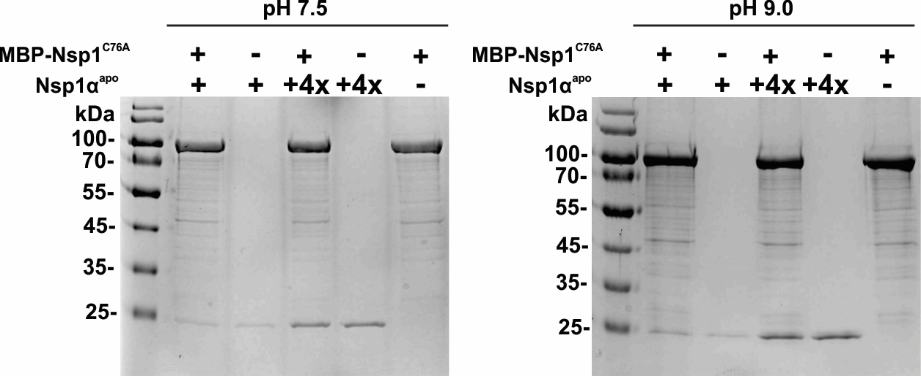

**Figure S8.** pH-dependent *in vitro* protease activity of WT Nsp1α^apo^, employing the full-length MBP-Nsp1^C76A^ protein as a substrate. Reactions contained 10 μM MBP-Nsp1^C76A^ and either 0.5 μM or 2 μM Nsp1α^apo^, incubated for 1 hr at 25 °C in 500 mM NaCl with either 50 mM HEPES, pH 7.5 (left panel), or 50 mM CHES, pH 9.0 (right panel). Cleavage products were resolved by SDS-PAGE and visualized by Coomassie staining.

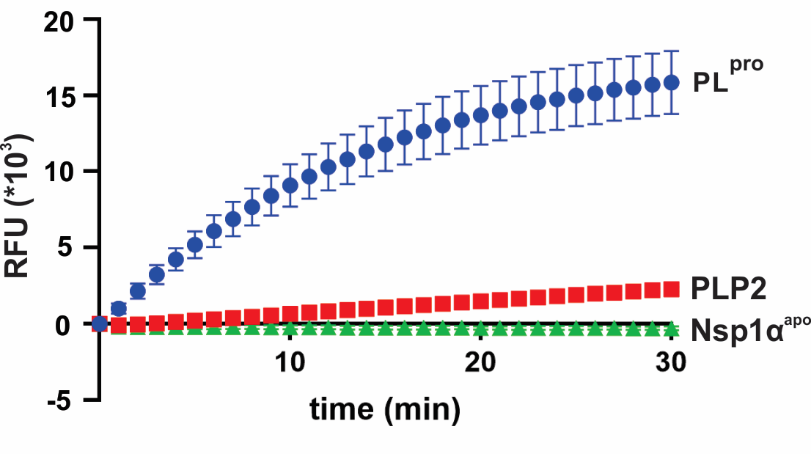

**Figure S9. Deubiquitinase activity assay**. Fluorescence-based deubiquitinase assay of Nsp1α^apo^, PL^pro^ from SAR-CoV, and PLP2 from PRRSV against Z-RLRGG-AMC. 1 μM enzyme was reacted with 25 μM peptide substrate. 96 well plates were preheated and maintained at 25 °C, and fluorescence was recorded after excitation at 340 nm (20 nm bandwidth) and emission was detected at 450 nm (20 nm). All fluorescence assays were performed in triplicate and data are shown as the mean ± the standard deviation (SD).

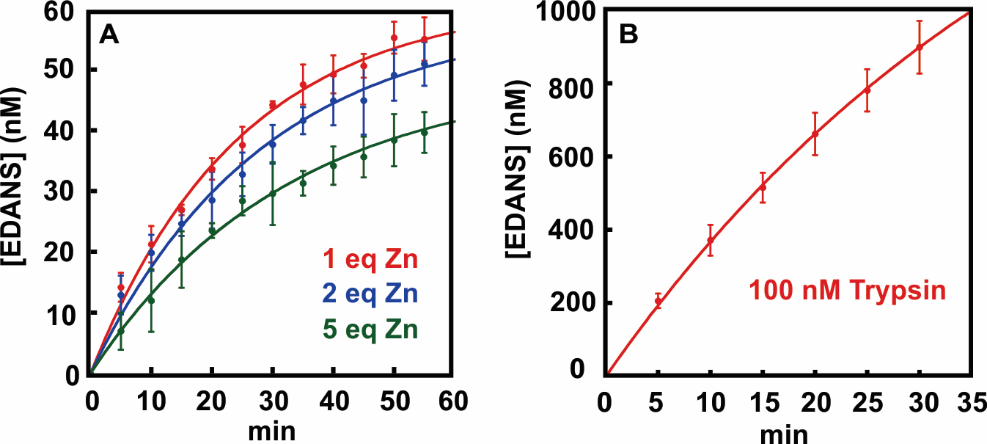

**Figure S10.** Fluorescence-based protease assay of Nsp1α^apo^ and trypsin against a fluorogenic peptide substrate containing the cleavage recognition sequence of Nsp1α (Dabcyl-FECAMATVYD-EDANS). (A) 20 μM Nsp1α^apo^ was reacted with 5 μM peptide substrate in the presence of 1 (red trace), 2 (blue trace), or 5 (green trace) molar eq of ZnCl_2_. (B) 0.1 μM trypsin was reacted with 5 μM peptide substrate (Dabcyl-FECAMATVYD-EDANS). The plates were preheated and maintained at 25 °C, and fluorescence was recorded after excitation at 360 nm (20 nm bandwidth) and emission was detected at 500 nm (20 nm). All fluorescence assays were performed in triplicates. Data are shown as the mean ± the standard deviation (SD).

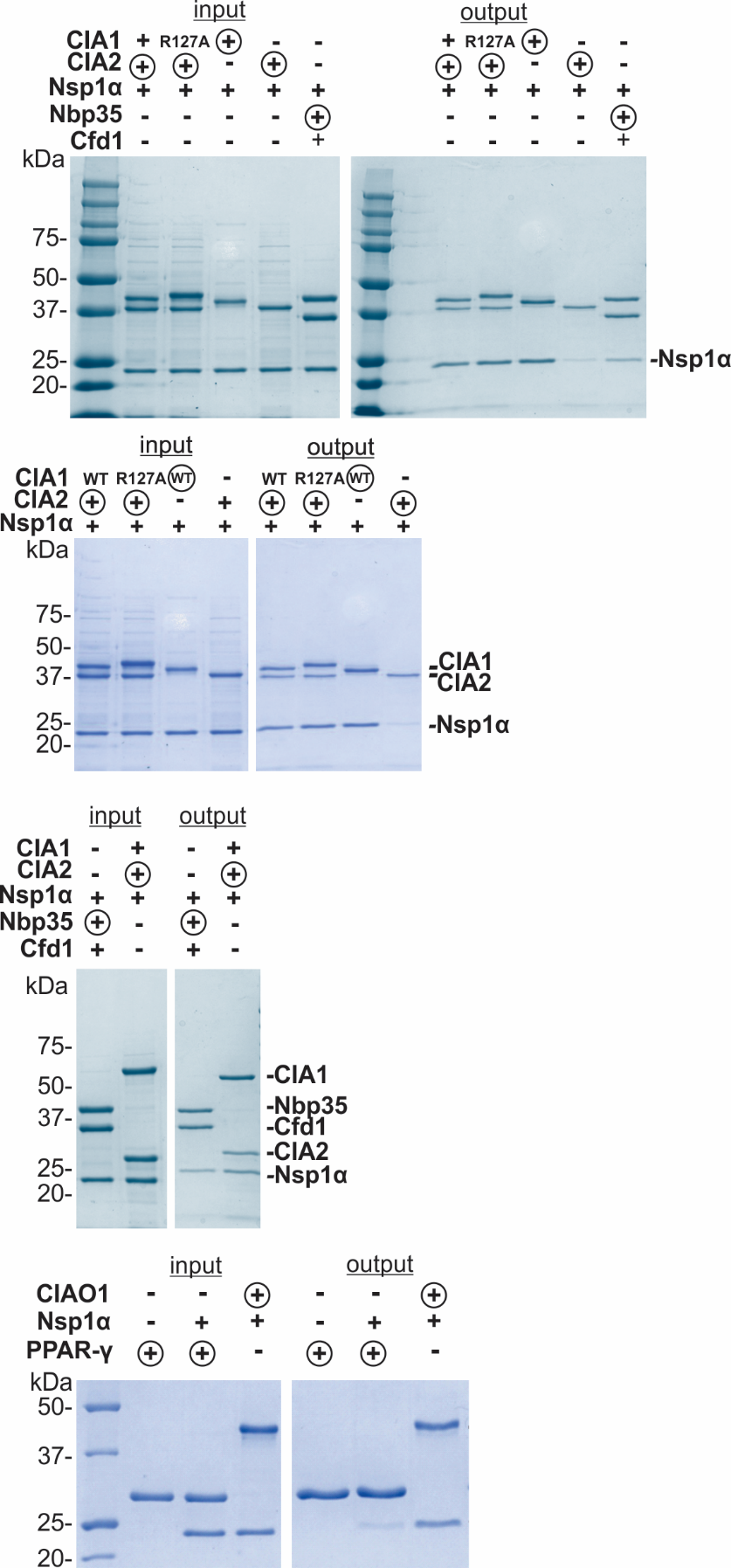

**Figure S11.** Pull-down assays assessing Nsp1α interactions with components of the *Chaetomium thermophilum* CTC (*Ct*CTC) via Strep-Tactin chromatography and monitored by SDS-PAGE. Circles indicate the immobilized “bait” protein carrying a Strep-Tag(II) affinity tag. Reactions were performed with 8–10 μM Strep-tagged bait protein and equimolar prey proteins, incubated in assay buffer (50 mM Tris, pH 8.0, 100 mM NaCl, 5 % glycerol, 5 mM β-mercaptoethanol) for 1 hr at 4 °C. Complexes were captured on Strep-Tactin XT Superflow resin, washed, and eluted with buffer containing 50 mM D-biotin and 100 mM NaOH. Elution fractions were resolved by SDS-PAGE (12 %) and visualized by Coomassie staining. The positions of the individual proteins are indicated on the side.

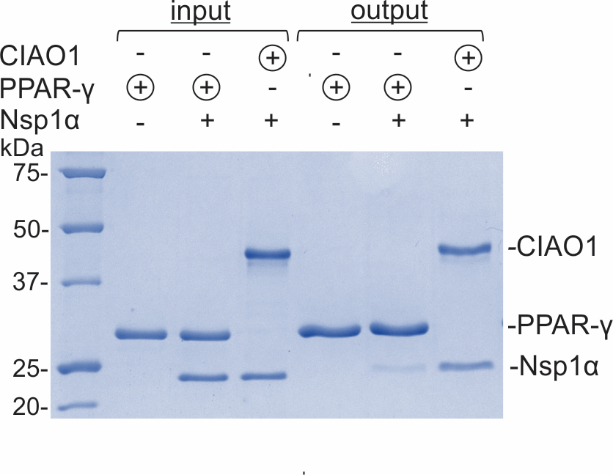

**Figure S12.** Pull-down assays assessing nonspecific binding of Nsp1α using PPAR-γ as a non-CIA bait control via Strep-Tactin chromatography and monitored by SDS-PAGE. Reactions were performed with 8–10 μM Strep-tagged bait protein (PPAR-γ or CIAO1) and equimolar Nsp1α, incubated in assay buffer (50 mM Tris, pH 8.0, 100 mM NaCl, 5 % glycerol, 5 mM β-mercaptoethanol) for 1 hr at 4 °C. Complexes were captured on Strep-Tactin XT Superflow resin, washed, and eluted with buffer containing 50 mM D-biotin and 100 mM NaOH. Elution fractions were analyzed by SDS–PAGE (12 %) and Coomassie staining.

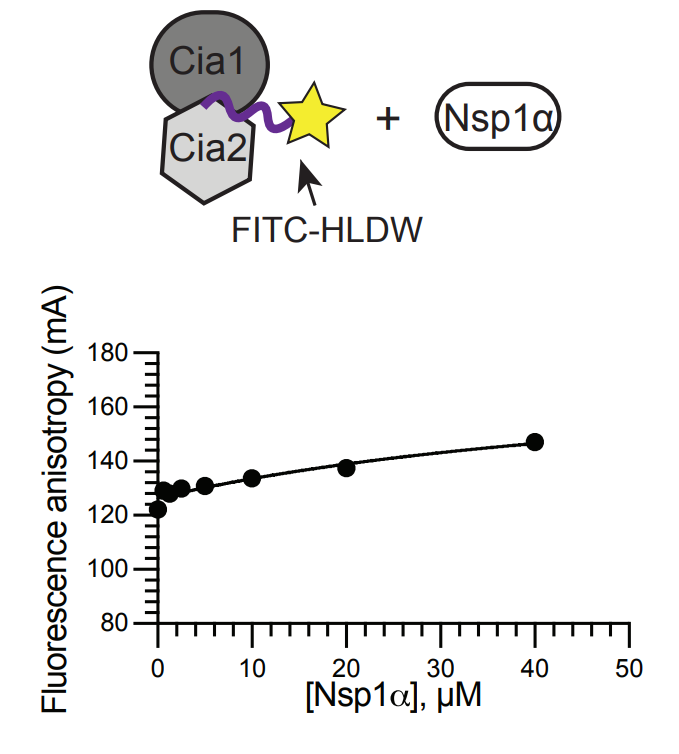

**Figure S13.** Competitive fluorescence anisotropy (FA) assay in which the preformed complex of FITC-HLDW (0.05 μM) with *Ct* Cia1-Cia2 (5 μM) was titrated with Nsp1α (0-40 μM). No decrease in anisotropy within the tested range indicates that Nsp1α does not bind the TCR binding site of the Cia1-Cia2 complex under the assay conditions.

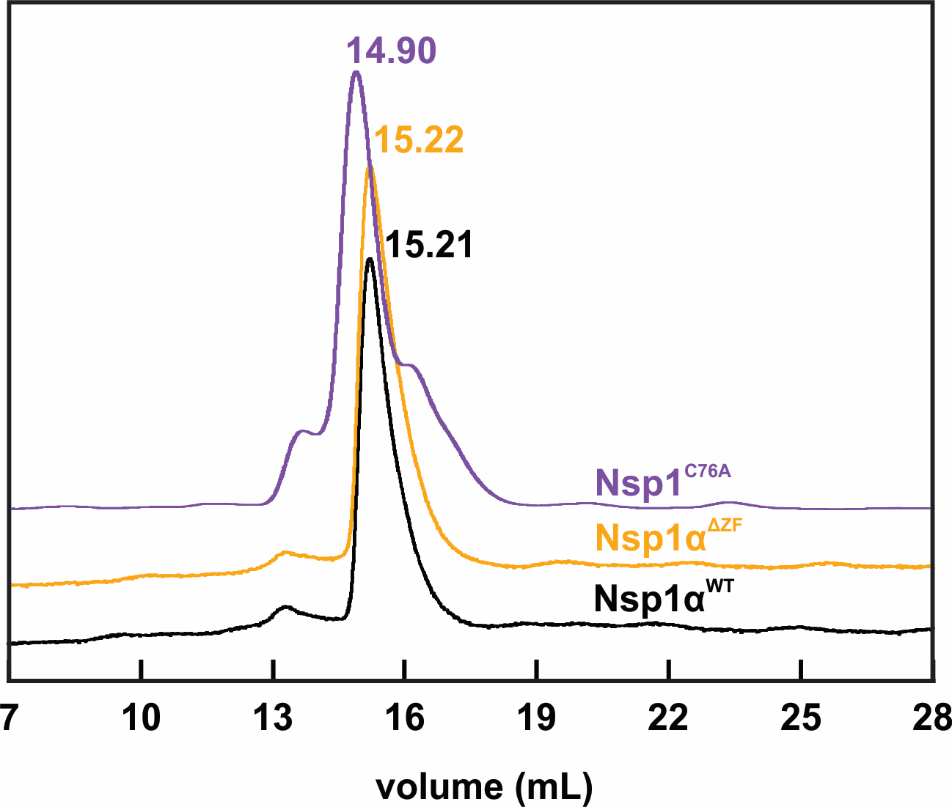

**Figure S14.** Analytical size-exclusion chromatography (SEC) of Nsp1α constructs*.* Traces for Nsp1α^WT^ (black), Nsp1α^ΔZF^ (orange), and Nsp1^C76A^ (purple) collected on a Superdex 200 Increase 10/300 GL column at 4 °C, 0.25 mL min⁻¹, with 100 µL injections of 50 µM protein. Running buffer: 50 mM HEPES, 500 mM NaCl, 0.5 mM TCEP, pH 8.0. The UV absorbance was monitored at 280 nm. Peak elution volumes (mL) are indicated (WT: 15.21, ΔZF: 15.22, C76A: 14.90). Traces are representative of n = 3 runs.

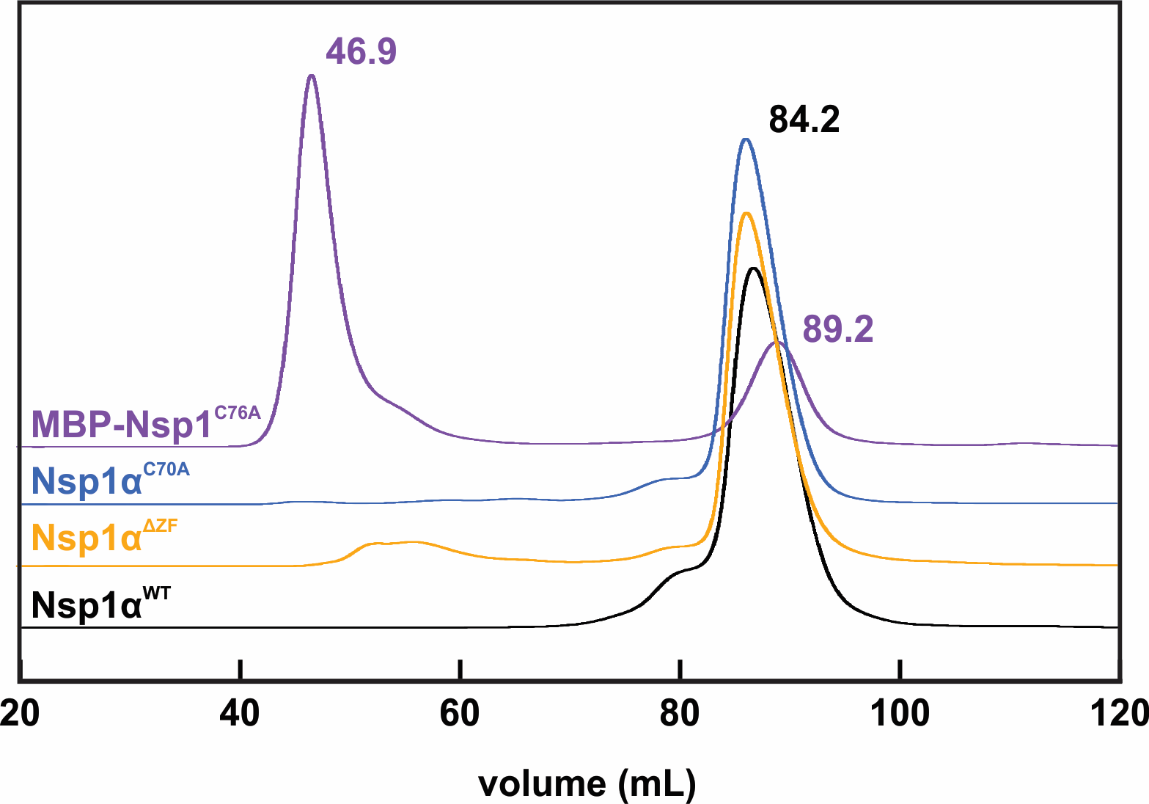

**Figure S15.** Preparative size-exclusion chromatography (SEC) of Nsp1α constructs. Traces for MBP–Nsp1^C76A^ (purple), Nsp1α^C70A^ (blue), Nsp1α^ΔZF^ (orange), and Nsp1α^WT^ (black) were collected on a Superdex 200 Increase 16/600 prep-grade column (Cytiva) at 0.5 mL min⁻¹. Running buffer: 50 mM HEPES, 500 mM NaCl, 0.5 mM TCEP, pH 8.0. Injections contained 1-5 mg of protein in 1 mL. UV absorbance was monitored at 280 nm. Elution volumes are indicated (MBP–Nsp1^C76A^, 46.9 and 89.2 mL; Nsp1α^WT^, 84.2 mL; Nsp1α^ΔZF^, 84.2 mL; Nsp1α^C70A^, 84.2 mL). The earlier peak at 46.9 mL corresponds to MBP–Nsp1^C76A^ (high oligomer elutes in the void volume of the column), whereas the Nsp1α constructs elute at 84 mL, consistent with their dimeric state. The peak at 89.2 mL corresponds to free MBP. Traces are representative of n = 3 runs.
